# Molecular features driving condensate formation and gene expression by the BRD4-NUT fusion oncoprotein are overlapping but distinct

**DOI:** 10.1101/2023.05.11.540414

**Authors:** Martyna Kosno, Simon L. Currie, Ashwani Kumar, Chao Xing, Michael K. Rosen

## Abstract

Aberrant formation of biomolecular condensates has been proposed to play a role in several cancers. The oncogenic fusion protein BRD4-NUT forms condensates and drives changes in gene expression in Nut Carcinoma (NC). Here we sought to understand the molecular elements of BRD4-NUT and its associated histone acetyltransferase (HAT), p300, that promote these activities. We determined that a minimal fragment of NUT (MIN) in fusion with BRD4 is necessary and sufficient to bind p300 and form condensates. Furthermore, a BRD4-p300 fusion protein also forms condensates and drives gene expression similarly to BRD4-NUT(MIN), suggesting the p300 fusion may mimic certain features of BRD4-NUT. The intrinsically disordered regions, transcription factor-binding domains, and HAT activity of p300 all collectively contribute to condensate formation by BRD4-p300, suggesting that these elements might contribute to condensate formation by BRD4-NUT. Conversely, only the HAT activity of BRD4-p300 appears necessary to mimic the transcriptional profile of cells expressing BRD4-NUT. Our results suggest a model for condensate formation by the BRD4-NUT:p300 complex involving a combination of positive feedback and phase separation, and show that multiple overlapping, yet distinct, regions of p300 contribute to condensate formation and transcriptional regulation.

## Introduction

Aberrant condensate formation occurs in a variety of cancers [1], including pediatric AML [2, 3], Ewing sarcoma [4], and some lung cancers [5]. Many such cancers are characterized by genetic translocations, resulting in expression of fusion oncoproteins, such as NUP98 fusions, EML4-ALK or EWS-FLI1 [1]. A number of these fusion proteins form condensates when expressed in cells through multivalent assembly of intrinsically disordered regions (IDRs) or modular signaling domains [2–5]. Condensate formation has been associated with the abnormal transcription or signaling that causes disease [2] although the link between condensates and transcription is not always clear [6]. The molecular mechanisms that cause condensate formation and might drive transcription or signaling by fusion oncoproteins are important in understanding cellular mechanisms of disease.

Aberrant biomolecular condensates form in Nut Carcinoma (NC), an aggressive, rare, and poorly differentiated cancer [7, 8]. On a molecular level the disease is characterized by chromosomal translocations resulting in a fusion protein that involves NUT and a partner protein. In most cases the partner is the transcriptional regulator, BRD4; the BRD4 homologue, BRD3; or the BRD4 ligand, NSD3 [9, 10]. The relationships between the NUT fusion partners suggest that the molecular mechanisms of the disease might be related among most patients.

BRD4 is a transcriptional co-activator that localizes at promotor and enhancer regions of the genome to stimulate gene expression [11]. The protein is composed of two bromodomains (BDs), an extra-terminal (ET) domain and a C-terminal intrinsically disordered region (IDR). The two BDs of BRD4 bind to acetylated histone tails, which promote its recruitment to acetylated, active chromatin. The ET domain interacts with additional elements of transcriptional machinery, including NSD3, JMJD6, CHD4, ATAD5 and GLTSCR1 [12, 13]. The C-terminal IDR similarly recruits factors such as the Mediator complex [14]. The collection of domains enables BRD4 to promote transcription initiation through assembly of these factors, which collectively recruit and activate RNA polymerase II to produce mRNA [12, 13].

NUT is normally expressed exclusively in male testes [9], where it plays an important role in spermatogenesis [15]. NUT is predicted to be mostly disordered throughout its 1150 amino acids, but contains a region, residues 355-505, that is predicted to be rich in α-helical structure. This element directly binds the histone acetyltransferase (HAT) protein, p300 [16, 17]. The interaction is mediated by two transactivation domains (TADs): TAD1 spanning residues 403-418 and TAD2 spanning residues 419-470 [18], which both bind the TAZ2 domain of p300. The TAD-TAZ2 interactions alleviate autoinhibition of the HAT domain by the TAZ2 domain, resulting in *trans*-autoacetylation of p300, which stimulates activity further [18]. In spermatogenesis, NUT binding to p300 orchestrates histone hyperacetylation, necessary for recruitment of a BRD4 homologue, BRDT [19, 20], leading to histone-to-protamine replacement [15].

It was previously shown that BRD4 can form intranuclear condensates, which are associated with super enhancers, to control gene expression [14]. The BRD4-NUT fusion protein also forms condensates in cells [21–23], which can contain over 2 Mbp of the genome. These condensates have been proposed to result from an aberrant positive feedback mechanism unique to the fusion protein [21, 22]. In this mechanism, BRD4-NUT at certain loci recruits and activates p300 through TAD-TAZ2 interactions. Activated p300 then produces high local levels of histone acetylation, driving recruitment of additional BRD4-NUT molecules, which recruit more p300 molecules, increasing acetylation further. This mechanism of condensate formation is different from that proposed for other fusion oncoproteins, e.g. NUP98 fusions, EML4-ALK or EWS-FLI1 [2–5], where multivalency-induced liquid-liquid phase separation (LLPS) has been invoked as the key driver. It is not clear whether LLPS may also contribute to condensate formation by BRD4-NUT. Since high levels of acetylation are often correlated with transcriptional activation, it is reasonable to suspect that condensate formation by BRD4-NUT, which involves p300-mediated acetylation, could be correlated with transcriptional changes in cells [23]. But it is not clear whether BRD4-NUT condensate formation is a significant driver of transcriptional changes.

Here, we sought to identify the molecular elements of NUT and p300 that promote formation of BRD4-NUT condensates and modulate gene expression. We developed a series of stable cell lines inducibly expressing different fusions of BRD4 and examined their capacity to form condensates and alter transcription. We found that the p300-interaction motif of NUT is necessary and sufficient, in fusion with BRD4, to form large condensates, and that condensate formation requires histone acetyl-transferase activity of p300. An engineered BRD4-p300 fusion protein induces condensates and gene expression similarly to BRD4-NUT, thus mimicking key aspects of the BRD4-NUT:p300 complex. With the BRD4-p300 fusion, condensate formation and transcriptional changes were partially distinguishable; p300 HAT activity is critical for transcriptional changes, whereas multiple molecular features of p300, including the HAT domain and an element predicted to undergo LLPS, collectively contribute to condensate formation. The data suggest a tentative model in which BRD4-NUT in complex with p300 forms condensates through a combination of positive feedback and LLPS, and that gene expression changes are driven partly by HAT activity of p300.

## Results

### BRD4-NUT forms condensates and recruits p300

We examined condensate formation and gene expression in a series of doxycycline-inducible 293TRex-FlpIn-based stable cell lines. To produce expression levels in the 293TRex-FlpIn that are similar to those in the HCC2429 patient-derived NC cell line, we extensively optimized doxycycline treatment and washout regimes (see methods and Fig.S1 and S2 for details). Once an optimal protocol was in hand, we first compared cells expressing the BRD4-NUT fusion protein, or BRD4 or NUT alone, each with an N-terminal mNeonGreen (Fig. 1a). All three proteins can form condensates in the 293TRex-FlpIn cells, observable by immunofluorescence (Fig. 1b) or mNeonGreen fluorescence (Fig. 1c). However, condensates formed by the BRD4-NUT fusion are appreciably larger and brighter than those formed by BRD4 or NUT (Fig. 1b). Furthermore, the percentage of cells forming large condensates (>1.25 µm in diameter, see Methods) is substantially higher in the BRD4-NUT line than in the two other lines (Fig. 1d).

**Fig. 1:**
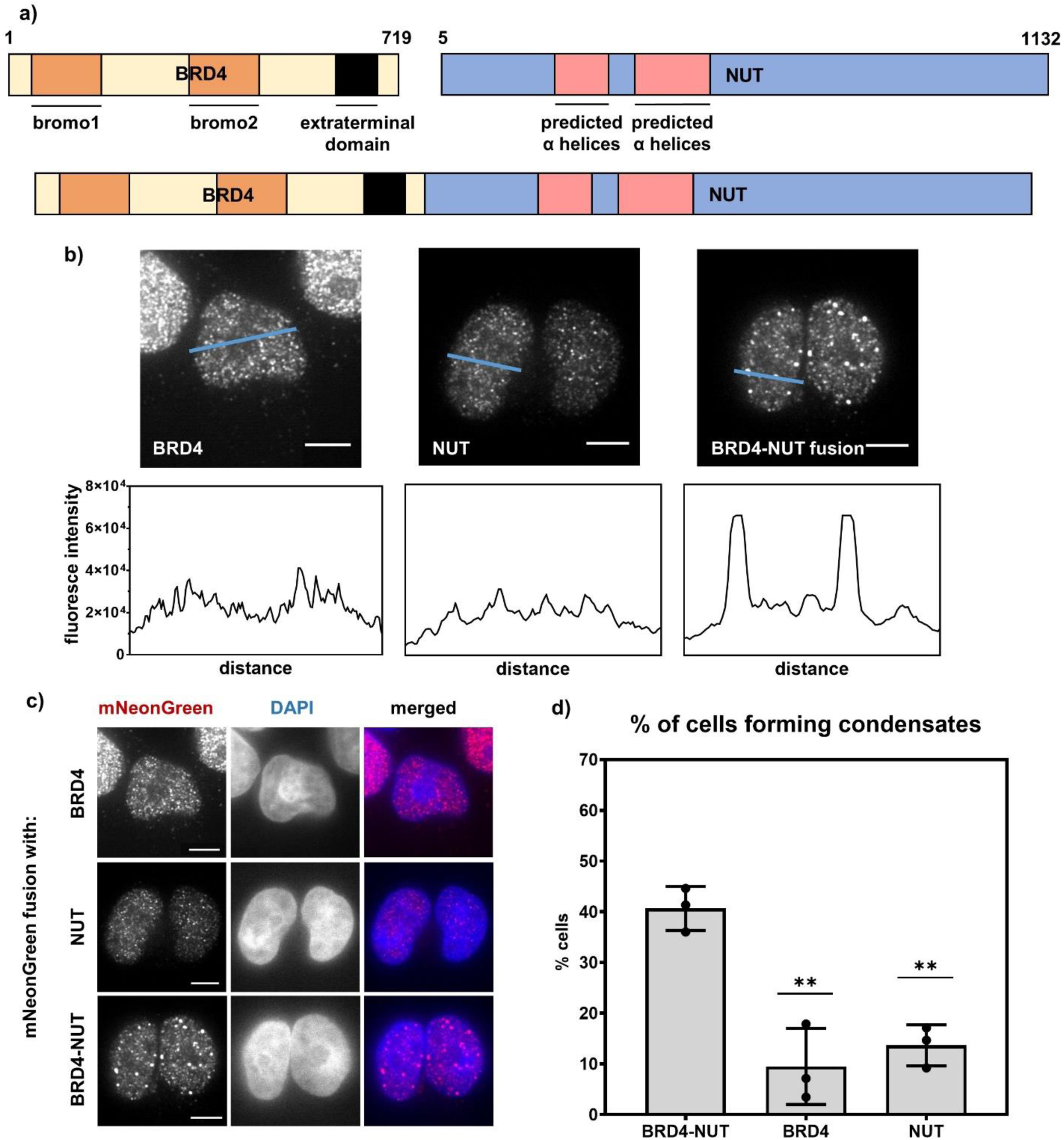
BRD4-NUT fusion protein forms nuclear condensates in human cells. a) Cartoon of BRD4, NUT and BRD4-NUT fusion proteins. Bromodomains 1 and 2 and extraterminal domain of BRD4 as well as predicted α-helices are indicated. b) Line profiles across cells expressing mNeonGreen-tagged BRD4, NUT and BRD4-NUT fusion; scale bars = 10μm. c) Micrographs of cells expressing mNeonGreen-tagged BRD4, NUT and BRD4-NUT; scale bars = 10μm. d) Quantification of percentage of cells forming condensates (at least 2 condensates larger than 1.25μm in diameter).

We next sought to confirm the recruitment of p300 into BRD4-NUT condensates (Fig. 2a) [16–18]. Pairwise co-immunostaining of patient-derived NC cells, HCC2429, for p300, BRD4 and NUT revealed colocalization of p300 with both other proteins/fragments, suggesting that p300 is recruited into BRD4-NUT condensates (Fig. 2b). Additionally, immunoprecipitation of BRD4-NUT from the stable cell line shows interaction with p300 (Fig. 2c), consistent with previously published data [16–18]. Together, these data indicate that the BRD4-NUT fusion protein has greater capacity to form condensates than either BRD4 or NUT alone, and that these condensates recruit p300, likely due to interaction of the fusion with p300 (see below).

**Fig. 2:**
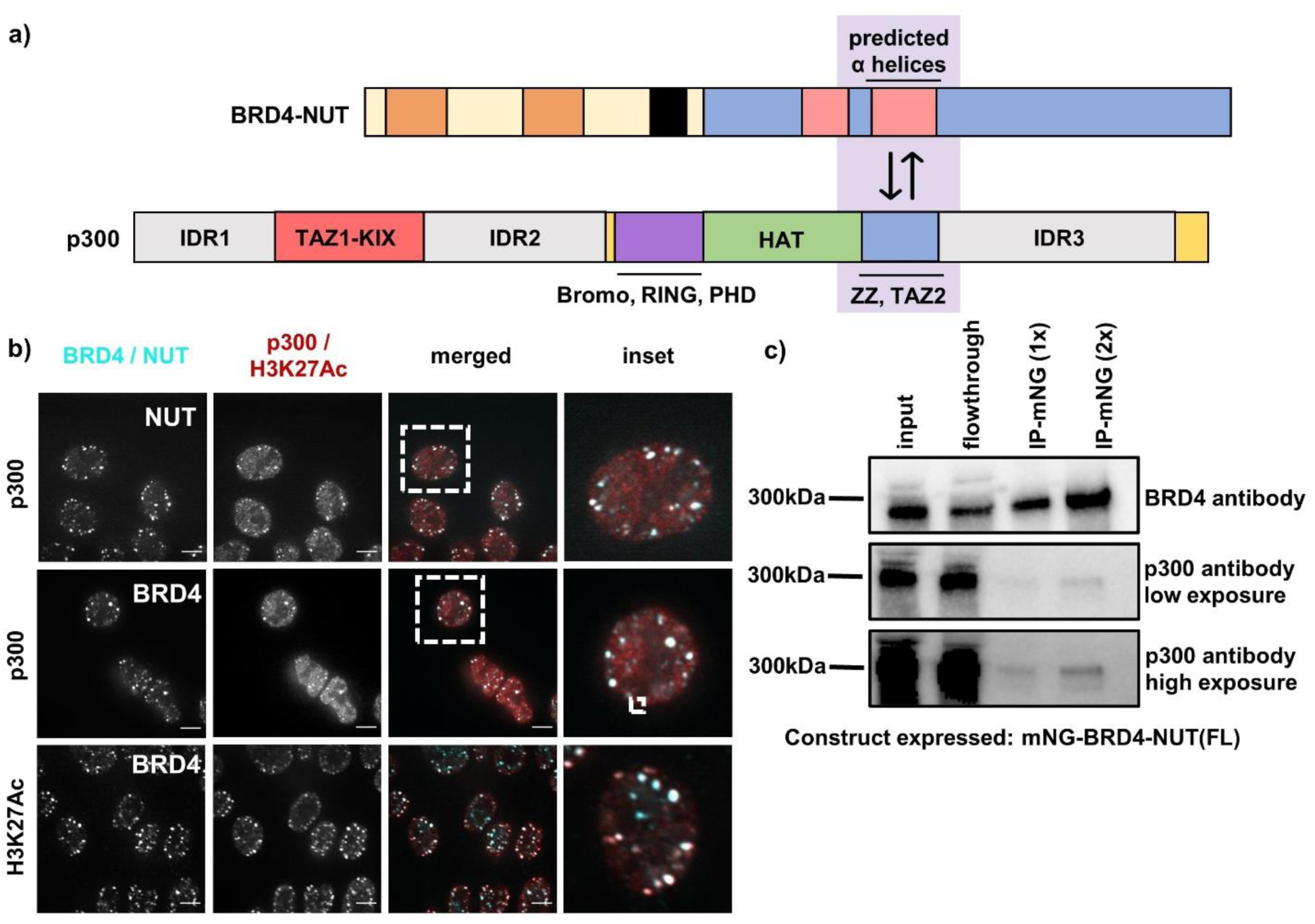
BRD4-NUT condensates recruit p300 histone acetyltransferase and are heavily acetylated at H3K27. a) Cartoon of BRD4-NUT and p300 interaction; known interaction motifs are indicated *(Shiota et al., Cell Rep., 2018; Reynoird et al., EMBO, 2010; Ibrahim et al., Nat Comm, 2022)* b) Micrographs of HCC2429 Nut Carcinoma cells co-stained with BRD4 and p300 or BRD4 and Histone H3K27Ac antibodies; scale bar = 10μm. c) Immunoprecipitation against mNeonGreen; western blot with a p300 antibody showing that BRD4-NUT pulls down p300.

### BRD4-NUT/p300 interaction is necessary and sufficient for condensate formation

We next questioned which part of NUT is required for condensate formation by BRD4-NUT. Residues 208-476 of NUT are predicted to form a series of α-helices (Fig.S3). An overlapping region of NUT, residues 346-593, has been shown to bind to p300 [16]. Very recently, crystal structures were reported of each of the two TADs of NUT in complex with the TAZ2 domain of p300 [18]. To preserve the structural elements within the p300-binding region, we designed a minimal fusion protein containing residues 355-505 of NUT, fused to BRD4, BRD4-NUT(MIN) and a complementary, BRD4-NUT(ΔMIN) fusion, lacking the MIN fragment of NUT (Fig. 3a). We then developed 293TRex-FlpIn-based stable cell lines inducibly expressing these constructs (Fig. 3b). While virtually no cells expressing BRD4-NUT(ΔMIN) contained large condensates, a similar fraction of cells expressing BRD4-NUT(MIN) or BRD4-NUT contained such structures (Fig. 3b,c). Like condensates produced by BRD4-NUT, those produced by BRD4-NUT(MIN) also recruit p300, as indicated by pairwise co-immunostaining of BRD4 and p300, and NUT and p300 (Fig. 3d,e). Conversely, mNeonGreen and p300 do not colocalize in cells expressing BRD4-NUT(ΔMIN) (Fig. 3f). Note that we used the α-mNeonGreen antibody to stain BRD4-NUT(ΔMIN) in these experiments because the α-NUT antibody fails to interact with this construct (Fig.S2c). We confirmed a high degree of colocalization between α-mNeonGreen and α-NUT antibodies in cells expressing BRD4-NUT, indicating that both antibodies recognize mNeonGreen-labeled BRD4-NUT construct to a similar extent (Fig.S1c). These data confirm that NUT interacts with p300 through its MIN fragment and suggest that this interaction is necessary and sufficient for condensate formation.

**Fig. 3:**
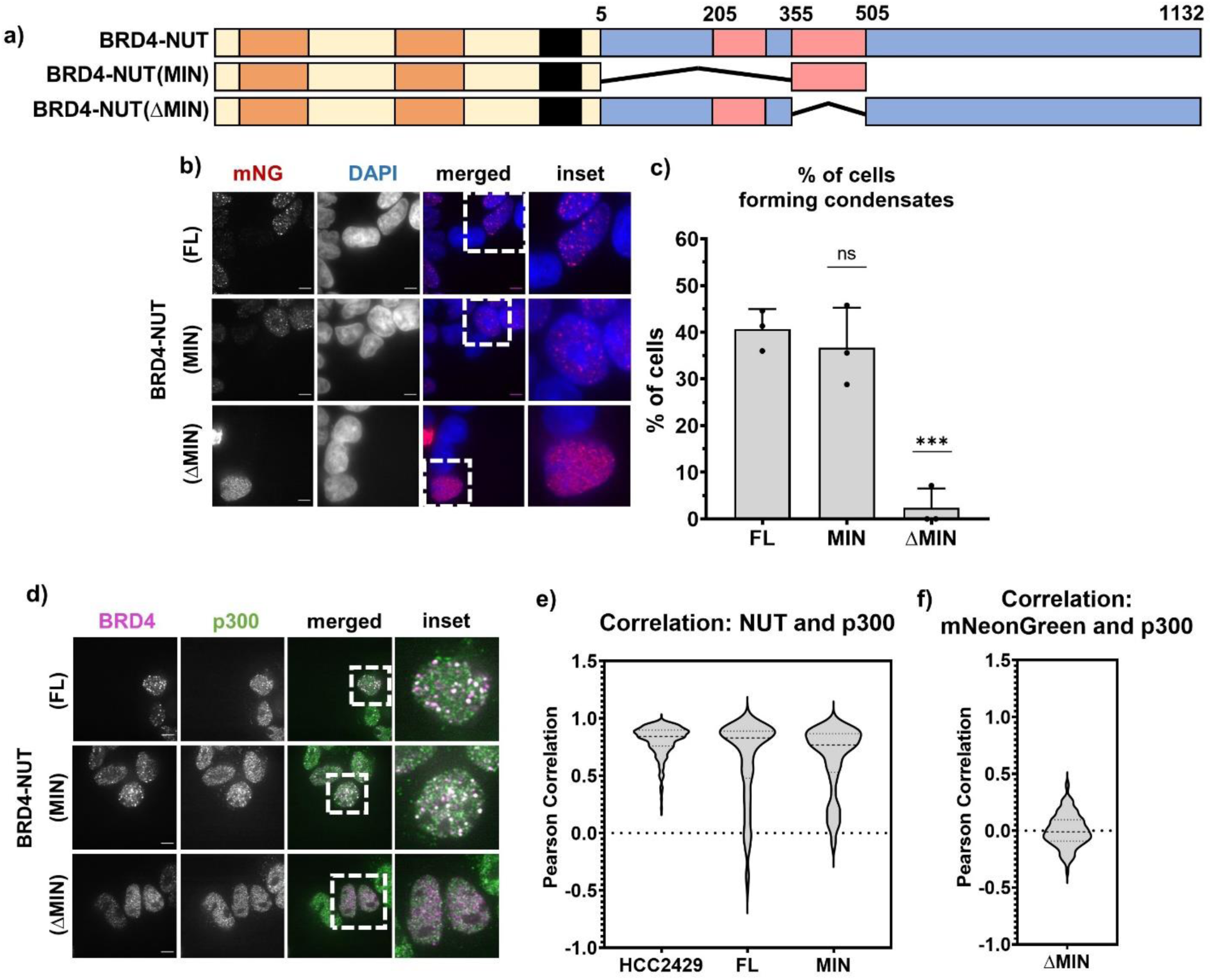
Minimal p300-interaction fragment of NUT in BRD4-NUT fusion is necessary and sufficient for condensate formation. a) Schematic of all BRD4-NUT – based constructs. b) Representative micrographs showing condensate formation in stable cell lines expressing different constructs. Staining against mNeonGreen shown in red, overlay with DAPI to indicate the nucleus. Scale bars = 10μm. c) Quantification of the micrographs represented in b. d) Representative micrographs comparing expression of constructs with or without p300-interaction motif in cells. Cells stained with anti-BRD4 antibody (magenta) and anti-p300 antibody (green). Scale bars = 10μm. e) and f) Quantification of the overlap between condensates via co-staining shown as Pearson correlation. Cells co-stained with NUT and p300 antibodies (left) or mNeonGreen and p300 antibodies (right); Nut antibody epitope was removed in BRD4-NUT(ΔMIN), see fig.S2c for more details.

Having established that p300 is recruited into BRD4-NUT condensates and that interaction of NUT with p300 is necessary to form condensates, we next asked whether p300 activity contributes to condensate formation. We treated both HCC2429 cells and the BRD4-fusion cell lines with JQ1, an inhibitor of bromodomain binding to acetylated histone tails [24], or with C646, an inhibitor of p300 histone acetyltransferase activity [25], and quantified the percentage of cells that form condensates. In all cases, C646 caused significant decreases in the fraction of cells containing large condensates (Fig. 4a, yellow bars). JQ1 caused even larger decreases for HCC2429 and the BRD4-NUT cell line but had a smaller effect on the BRD4-NUT(MIN) cells that was not statistically significant, perhaps due to a larger variability in the data (Fig. 4a, pink bars). These data suggest that both histone acetylation and BRD4 binding to acetylated histones contribute to BRD4-NUT condensate formation.

**Fig. 4:**
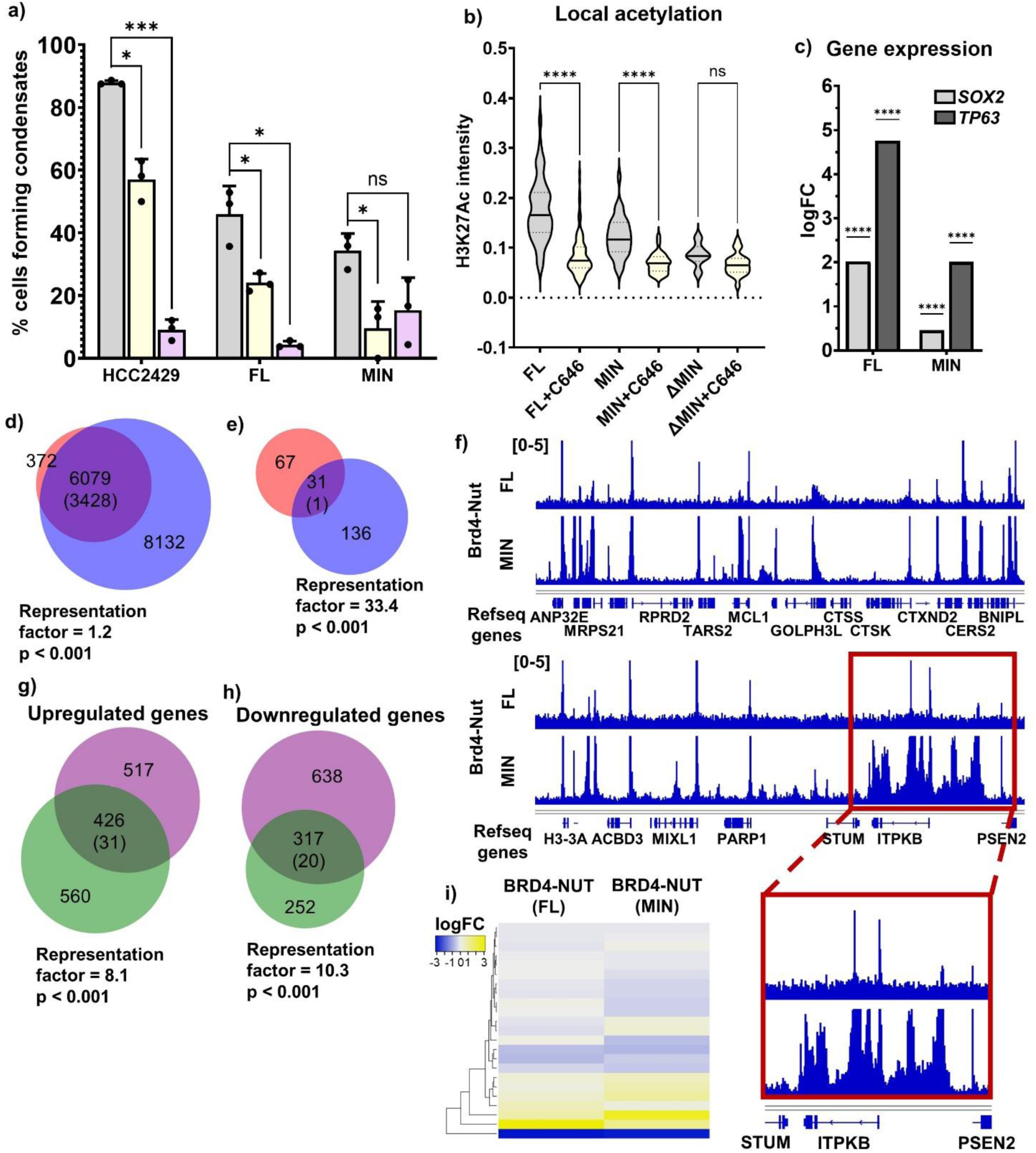
p300 binding and enzymatic activity are necessary for gene expression changes induced by BRD4-NUT. a) Quantification of cells capacity to form condensates, comparing cells that are untreated (gray) and cells treated with C646 inhibitor (yellow) or JQ1 inhibitor (pink). Cells compared here include HCC2429 Nut Carcinoma cell line and stable cell lines expressing BRD4-NUT(FL) and BRD4-NUT(MIN). b) Local acetylation measured as average fluorescence intensity across a condensate. Data shown for cells untreated (gray) or treated with C646 (yellow), in cells expressing BRD4-NUT(FL), BRD4-NUT(MIN) and BRD4-NUT(ΔMIN). c) Graph showing logFC for *SOX2* and *TP63* genes upon expression of different BRD4-p300 constructs. d) Venn diagram of gene occupancy by BRD4-NUT(FL) (pink) and BRD4-NUT(MIN) (blue), found via ChIPseq with @NUT antibody: all annotated genes occupied by either protein. Values in the diagram show the number of unique genes annotated; numbers in parentheses represent the mean overlap in 20 iterations with a randomly generated gene pool of the same size, from the human genome. e) Venn diagram of gene occupancy by BRD4-NUT(FL) (pink) and BRD4-NUT(MIN) (blue), found via ChIPseq with @NUT antibody: genes annotated at promoter-TSS regions of genes, with normalized signal value of 0.3-1. Values in the diagram as in d). f) Example ChIPseq tracks comparing gene occupancy by BRD4-NUT(FL) and BRD4-NUT(MIN). Data scale: 0-5. Inset shows more closely an example of the same genes being occupied by both proteins, but BRD4-NUT(MIN) occupying more loci. g) Venn diagram of genes upregulated upon expression of BRD4-NUT(FL) (purple) or BRD4-NUT(MIN) (green), found via RNAseq. Genes shown when fold change in expression was greater than two and p<0.05. Numbers in parentheses represent the mean overlap in 20 iterations with a randomly generated gene pool of the same size, from the human genome. i) Venn diagram of genes downregulated upon expression of BRD4-NUT(FL) (purple) or BRD4-NUT(MIN) (green), found via RNAseq. Statistical analyses as in g). j) RNAseq-ChIPseq data integration: heatmap showing up-and down-regulated genes found via RNAseq, bound by both BRD4-NUT(FL) and BRD4-NUT(MIN). Top 200 genes shown in the heatmap, based on p value.

Finally, we assessed p300 activity in BRD4-NUT(MIN) condensates through examining the histone post-translational modification, H3K27Ac, a well-established marker of active transcription that is introduced by p300 [26]. We used immunofluorescence to quantify the H3K27Ac mark within condensates via confocal microscopy and used it as a proxy for the localized acetylation level. We found that acetylation in BRD4-NUT and BRD4-NUT(MIN) is relatively high and decreases upon treatment with C646 (Fig. 4b). As described above, only a small fraction of cells expressing BRD4-NUT(ΔMIN) form large condensates (Fig. 3b,c). With the caveat of a restricted-size sample, we observed that acetylation within BRD4-NUT(ΔMIN) condensates was significantly lower than in BRD4-NUT or BRD4-NUT(MIN) condensates, and did not further decrease upon C646 treatment (Fig. 4b). These behaviors are consistent with the lack of co-localization between p300 and BRD4-NUT(ΔMIN) (Fig. 3f). Thus, p300 acetylates histones within BRD4-NUT condensates.

Together, these data show that binding of p300 to the helical motifs in NUT produces a BRD4-NUT:p300 complex in which HAT activity contributes to condensate formation and drives local histone acetylation.

### BRD4-NUT and BRD4-NUT(MIN) have highly similar transcriptional profiles

We next sought to examine the functional consequences of the interaction between BRD4-NUT and p300. Specifically, we studied the transcriptional changes that occur upon expression of BRD4-NUT and the potential role of p300 in driving these changes. We performed RNAseq and ChIPseq (with an α-NUT antibody) on the BRD4-NUT and BRD4-NUT(MIN) cell lines as well as the parent 293TRex-FlpIn line (RNAseq only), each in two biological replicates.

The BRD4-NUT cells show substantial differences in gene expression compared to the parental cells. Some of the differentially expressed genes have been reported as signature genes of NC, including *SOX2* and *TP63* [8, 21] (Fig. 4c). Additionally, RNAseq-based Ingenuity Pathway Analysis (IPA) revealed that expression of BRD4-NUT causes significant up-and downregulation of multiple cellular pathways, many of which are related to cancer development and progression, including: VDR and RXR activation, GP6 signaling, epithelial-mesenchymal transition, Netrin-1 signaling, WNT signaling, Basal Cell Carcinoma and Glioblastoma Multiforme signaling [27–31] (Fig.S4a). These changes in cellular pathways support that the expression of BRD4-NUT in 293TRex-FlpIn cells results in relevant gene expression changes.

To examine the importance of the MIN fragment of NUT, we compared genome occupancy and gene expression in the BRD4-NUT and BRD4-NUT(MIN) lines. The ChIPseq data reveal that most genes bound by BRD4-NUT are also bound by BRD4-NUT(MIN) (Fig. 4d). Furthermore, when the ChIPseq analysis is limited to only the promoter or transcription start site (TSS) regions, 31 genes are commonly bound by BRD4-NUT and BRD4-NUT(MIN), which is much higher than expected at random (1) (Fig. 4e), as indicated by p-value and representation factor (see Methods). While occupancy patterns are similar for the two proteins, many more genes are bound by BRD4-NUT(MIN) than by BRD4-NUT (Fig. 4d), suggesting that gene occupancy is more restricted for the full-length fusion. This finding was corroborated upon closer examination of ChIPseq tracks, with the majority of peaks coinciding at the same genomic loci for both cell lines, but some additional peaks only present with BRD4-NUT(MIN) (Fig. 4f).

Analogous to ChIPseq, the RNAseq data show a large portion of differentially-expressed genes in common between the BRD4-NUT and BRD4-NUT(MIN) cell lines (Fig. 4g,h). The number of overlapping genes in RNAseq is more than 10 times higher than expected at random, for both up-and downregulated genes, indicating a significant overlap in transcriptional profiles of the two cell lines (Fig. 4g,h). We also integrated the ChIPseq and RNAseq results for cells expressing either construct. Here, we focused on genes occupied by both proteins, as measured by ChIPseq annotations at promoter-TSS (Fig. 4e, intersection in the diagram). We then analyzed the expression of these genes via RNAseq and found that they are similarly up-or downregulated in both cell lines (Fig. 4i). Several of the commonly upregulated genes encode transcriptional regulators, including: CHD3, FOXM1 SETD2, SETD5 and MSL1, as well as proteins involved in apoptotic signaling, including: EIF5A, NGFRAP1 and FAIM. Notably, the upregulated gene *ZMYND11* encodes a zinc finger protein that is architecturally similar to ZMYND8, which is fused with NUT in some cases of NC [32]. These analyses suggest that BRD4-NUT and BRD4-NUT(MIN) both bind to similar genes and induce similar transcriptional changes. Thus, BRD4-NUT(MIN) is sufficient to recapitulate the majority of transcriptional changes caused by BRD4-NUT. This similarity suggests that the gene expression changes are due to either the BRD4 portion of the two proteins, or to the ability of BRD4-NUT to bind p300 through the MIN element (or both). The data below suggest that p300 binding plays an important role, but do not rule out effects from BRD4 also.

### BRD4-p300 recapitulates BRD4-NUT – mediated condensate formation and transcriptional changes

Since BRD4-NUT(MIN) mimics condensate formation and transcriptional activity of BRD4-NUT, we hypothesized that recruitment of p300 into BRD4-NUT condensates may be the main function of NUT sequences in the fusion protein. To study the roles of p300 and dissect its molecular features, we thus fused BRD4 directly to p300 (Fig. 5a). This approach eliminates any potential additional functions of NUT and focuses solely on its ability to recruit p300. We established a new 293TRex-FlpIn-based stable cell line, expressing an mNeonGreen-tagged BRD4-p300 fusion protein. The expression of this fusion is similar to that of BRD4-NUT (Fig.S2b), and the two lines form condensates to a similar degree (Fig. 5b,c). We further measured average local acetylation levels within condensates, via immunostaining with an α-H3K27Ac antibody. Stable cell lines expressing either BRD4-p300, BRD4-NUT or BRD4-NUT(MIN) all show elevated acetylation levels, which decrease upon inhibition of p300 with C646 (Fig. 5d). Thus, the BRD4-p300 construct causes condensate formation to a similar degree as BRD4-NUT and BRD4-NUT(MIN), and all three constructs cause elevated acetylation.

**Fig. 5:**
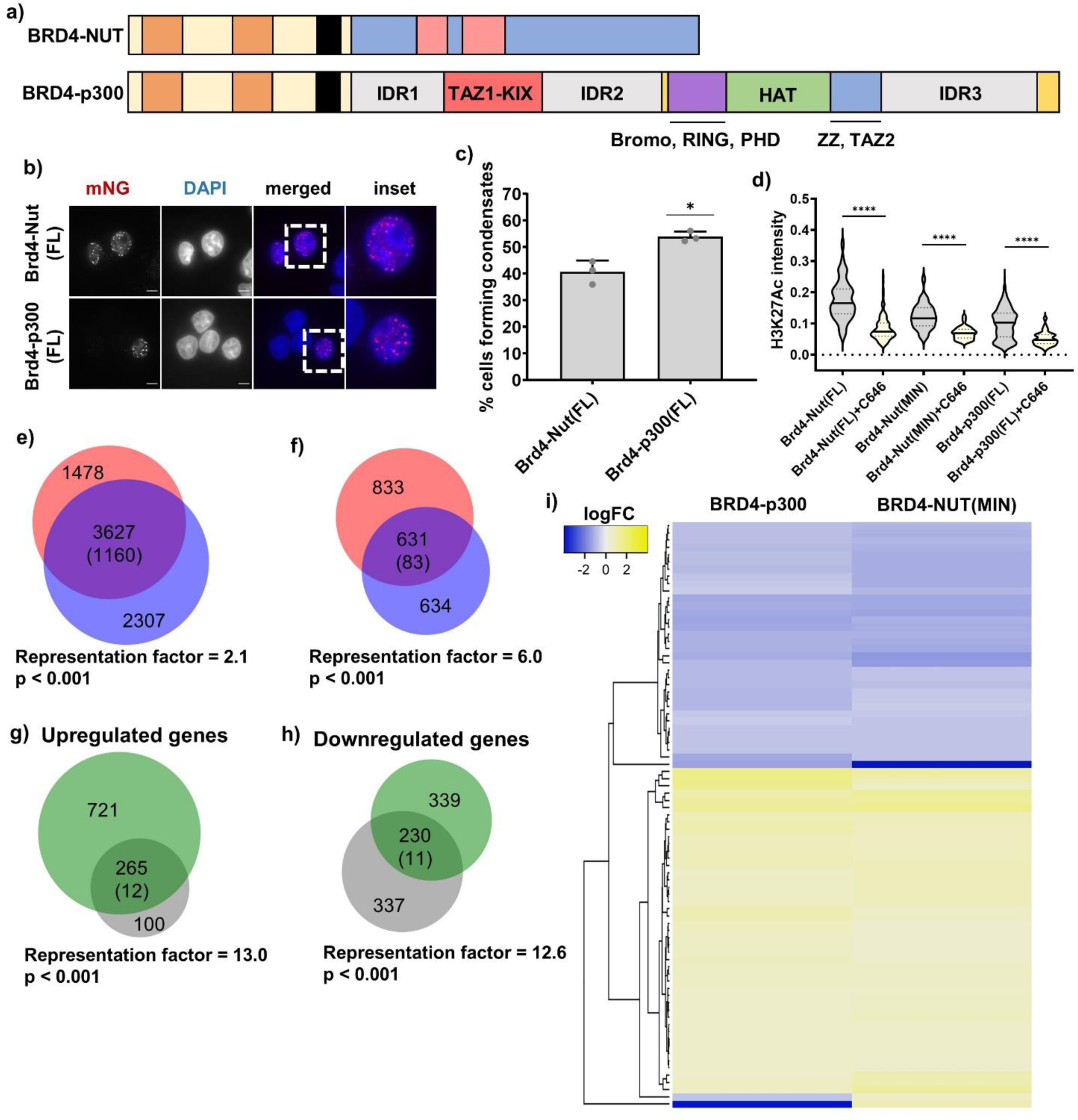
Brd4-p300 fusion forms condensates and recapitulates many of the transcriptional changes observed for Brd4-Nut fusion. a) Schematic of Brd4-Nut and Brd4-p300. b) Micrographs of Brd4-Nut and Brd4-p300 – expressing stable cell lines. Cells are stained with mNeonGreen antibody; scale bars = 10μm. c) Quantification of the micrographs represented in b. d) Quantification of histone H3K27Ac staining in untreated vs. C646-treated cells. Local acetylation shown as average fluorescence intensity per condensate. e) Venn diagram of genes acetylated upon expression of BRD4-NUT(MIN) (pink) and BRD4-p300 (blue), found via ChIPseq with @H3K27Ac antibody, at promoter-TSS regions: all annotated genes. Values in the diagram show the number of unique genes annotated; numbers in parentheses represent the mean overlap in 20 iterations with a randomly generated gene pool of the same size, from the human genome. f) Venn diagram of genes acetylated upon expression of BRD4-NUT(MIN) (pink) and BRD4-p300 (blue), found via ChIPseq with @H3K27Ac antibody, at promoter-TSS regions: Diagram shows only genes after applying 0.3-1 normalized signal value cutoff. Values in the diagram as in e). g) Venn diagram of genes upregulated upon expression of BRD4-NUT(MIN) (green) or BRD4-p300 (gray), found via RNAseq. Genes shown when fold change in expression was greater than two and p<0.05. Numbers in parentheses represent the mean overlap in 20 iterations with a randomly generated gene pool of the same size, from the human genome. h) Venn diagram of genes downregulated upon expression of BRD4-NUT(MIN) (green) or BRD4-p300 (gray), found via RNAseq. Statistical analyses as in g). i) RNAseq-ChIPseq data integration: heatmap showing up-and down-regulated genes found via RNAseq, that are bound by both BRD4-NUT(MIN) and BRD4-p300. Top up-and downregulated genes shown in the heatmap, based on log fold change, where log fold change was greater than 0.5.

Based on our data, we expected the BRD4-p300 construct to most closely resemble BRD4-NUT(MIN) in its activity, including condensate formation and transcriptional regulation. Due to the lack of a NUT antibody epitope in BRD4-p300, and the presence of wild-type BRD4 and p300 in the cellular background in all our stable cell lines, we could not perform a ChIPseq-based gene occupancy analysis. Instead, we examined genome-wide acetylation via ChIPseq against the H3K27Ac mark. We found that almost all of the same genes are acetylated on histone H3K27 upon expression of either BRD4-NUT(MIN) or BRD4-p300 (Fig. 5e). When we restrict the analysis to only gene annotations with the highest normalized signal value, overlap between the two cell lines is still significant, with the number of overlapping genes more than 7 times higher than predicted at random (Fig. 5f). Thus, BRD4-p300 and BRD4-NUT(MIN) bind similar genomic loci. We also analyzed RNAseq results from the stable cell lines and compared them to the control 293TRex-FlpIn cells. We found that a large fraction of differentially expressed genes are commonly up-or downregulated in both BRD4-NUT(MIN) and BRD4-p300 – expressing cells (Fig. 5g,h). The fraction of genes modulated in both cell lines is more than 20-fold greater than expected at random (Fig. 5g,h). Finally, we integrated ChIPseq and RNAseq data, by analyzing expression patterns of genes commonly acetylated upon expression of either BRD4-NUT(MIN) or BRD4-p300 (Fig. 5f, overlap in the diagram). Out of the 631 genes, we limited our analysis to 81 shown in the heatmap (Fig. 5i), based on the most significant log fold change. This analysis showed that indeed both cell lines present highly similar differential gene expression patterns (Fig. 5i). Many genes found through this analysis encode zinc finger transcription factors, including: *ZBTB25*, *ZNF213*, *ZNF644*, *ZNF408*, *ZNF583* and *ZMYND8.* As noted above, ZMYND8 was previously reported as a fusion partners of NUT in NC [32].

Thus, BRD4-NUT(MIN) and BRD4-p300 behave similarly in terms of driving condensate formation and transcriptional changes. These results support the importance of the MIN fragment of NUT in recruiting p300 to BRD4-NUT, and suggest that p300 is responsible for a significant portion of transcriptional changes observed in BRD4-NUT – expressing cells.

### p300 IDRs, TF binding and enzymatic activity contribute to condensate formation

Having shown that recruitment of p300, either through binding [BRD4-NUT(MIN)] or through covalent attachment (BRD4-p300), is sufficient to recapitulate condensate formation and transcriptional changes in cells expressing BRD4-NUT, we next sought to learn what molecular features of p300 are responsible for these behaviors. p300 contains many domains within its structure, falling into three classes of molecular features: (1) the histone acetyltransferase domain (HAT), here referred to as “H”, (2) transcription factor-binding domains (bromodomain, PHD domain, ZZ, TAZ1, TAZ2, KIX, and RING), here collectively named “T”, and (3) three predicted intrinsically disordered regions (IDRs: IDR1, IDR2 and IDR3) (Fig. 6a), collectively referred to as “I”. We designed a series of BRD4-p300 fusion mutants, with different portions of p300 deleted or inactivated.

**Fig. 6:**
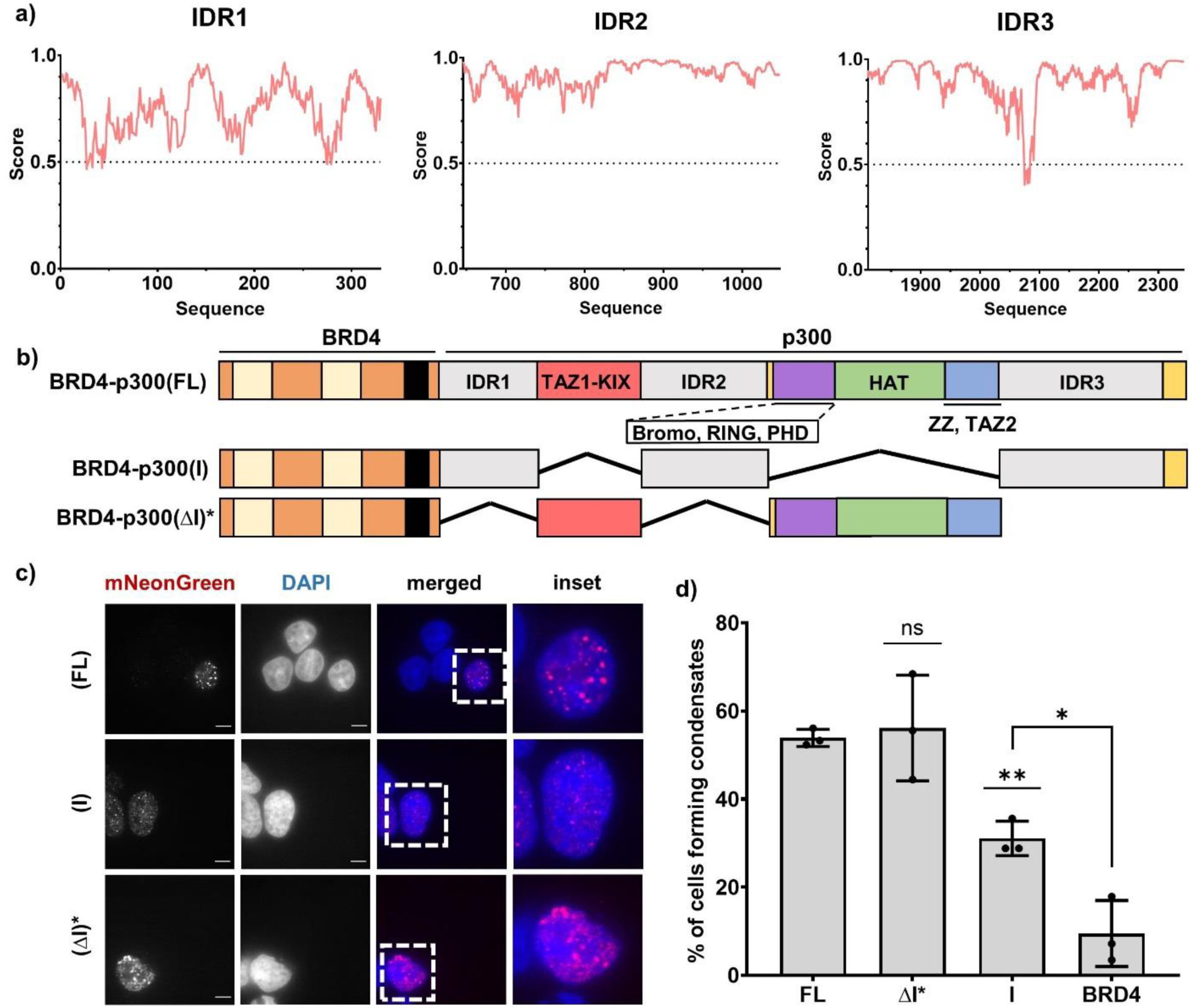
p300 IDRs are dispensable for condensate formation. a) Disorder prediction of p300 IDRs as shown via IUPRED2 analysis. b) Schematics of BRD4-p300(FL), (IDR) and (ΔIDR) constructs; different domains and disordered regions are indicated. c) Micrographs of all the BRD4-p300 constructs from a); scale bars = 10μm. d) Quantification of the micrographs represented in c).

Since protein IDRs are often important in formation of biomolecular condensates [33–37], we first focused on the roles of p300 IDRs, which are predicted to be disordered throughout their lengths by IUPRED [38, 39] (Fig. 6a). All three of the sequences are classified as “weak polyampholytes and polyelectrolytes” by CIDER [40] due to their low overall charge, suggesting a propensity to undergo LLPS. They are also predicted to phase separate by PSPredictor [41] (Fig.S5). Thus, the sequences of the IDRs indicate they likely can self-assemble and phase separate in physiologic conditions. We generated two new constructs based on the mNeonGreen-BRD4-p300 fusion: (1) IDRs of p300 fused to BRD4 [BRD4-p300(I)], and (2) IDRs deleted, with only the structured domains of p300 fused to BRD4 [BRD4-p300(ΔI)] (Fig. 6c). We attempted to develop both stable cell lines but were only successful with the BRD4-p300(I) construct; expression of the BRD4-p300(ΔI) in cells was toxic. Thus, all experiments with BRD4-p300(ΔI) were performed via transient transfections. To compare the BRD4-p300(ΔI) construct in transient transfections with other constructs in stable cell lines, we applied stringent expression level cutoffs (based on mNeonGreen fluorescence) to our single-cell image analyses (Fig.S2, S6). We could not apply a similar expression cutoff to pooled-cell transcriptional analyses; thus, we did not perform RNAseq or ChIPseq with the BRD4-p300(ΔI) construct.

In imaging analyses we found that a higher fraction of cells expressing BRD4-p300(I) contain large condensates than those expressing BRD4 alone, indicating that the IDRs in p300 contribute to condensate formation (Fig. 6d,e). However, fewer BRD4-p300(I) cells contain condensates than BRD4-p300 cells (Fig. 6d,e), indicating that other elements of p300 also are important in this regard. Somewhat surprisingly given the BRD4/BRD4-p300(I) comparison, cells expressing BRD4-p300(ΔI) form condensates to a similar extent as BRD4-p300 (Fig. 6d,e). However, the morphology of these condensates is different: those produced by BRD4-p300(ΔI) are larger and less round than those formed by BRD4-NUT (Fig. 6d, S4d), suggesting they may not be physically different and not quantitatively comparable. Thus, while the IDRs of p300 are not the sole driver of BRD4-p300 condensation, they promote condensate formation when fused to BRD4.

We next tested whether the structured domains of p300 are important in condensate formation. We designed two additional constructs: (1) BRD4-p300(H*IT), which has a HAT-inactivating point mutation (D1399Y [42]) but retains the IDRs and other structured domains and (2) BRD4-p300(HI), where the p300 IDRs and HAT domain are intact but the TF-binding domains are deleted (Fig. 7a). We found that inactivating the HAT domain via a point mutation in BRD4-p300(H*IT), or removing transcription factor-binding domains in BRD4-p300(HI), both decrease the fraction of cells forming condensates relative to BRD4-p300 (Fig. 7b,c). Both mutants, however, have higher condensate forming activity than BRD4 alone (Fig. 7b,c). These results suggest that all three classes of molecular features of p300—IDRs, HAT and TF-binding domains— contribute to formation of condensates in cells.

**Fig. 7:**
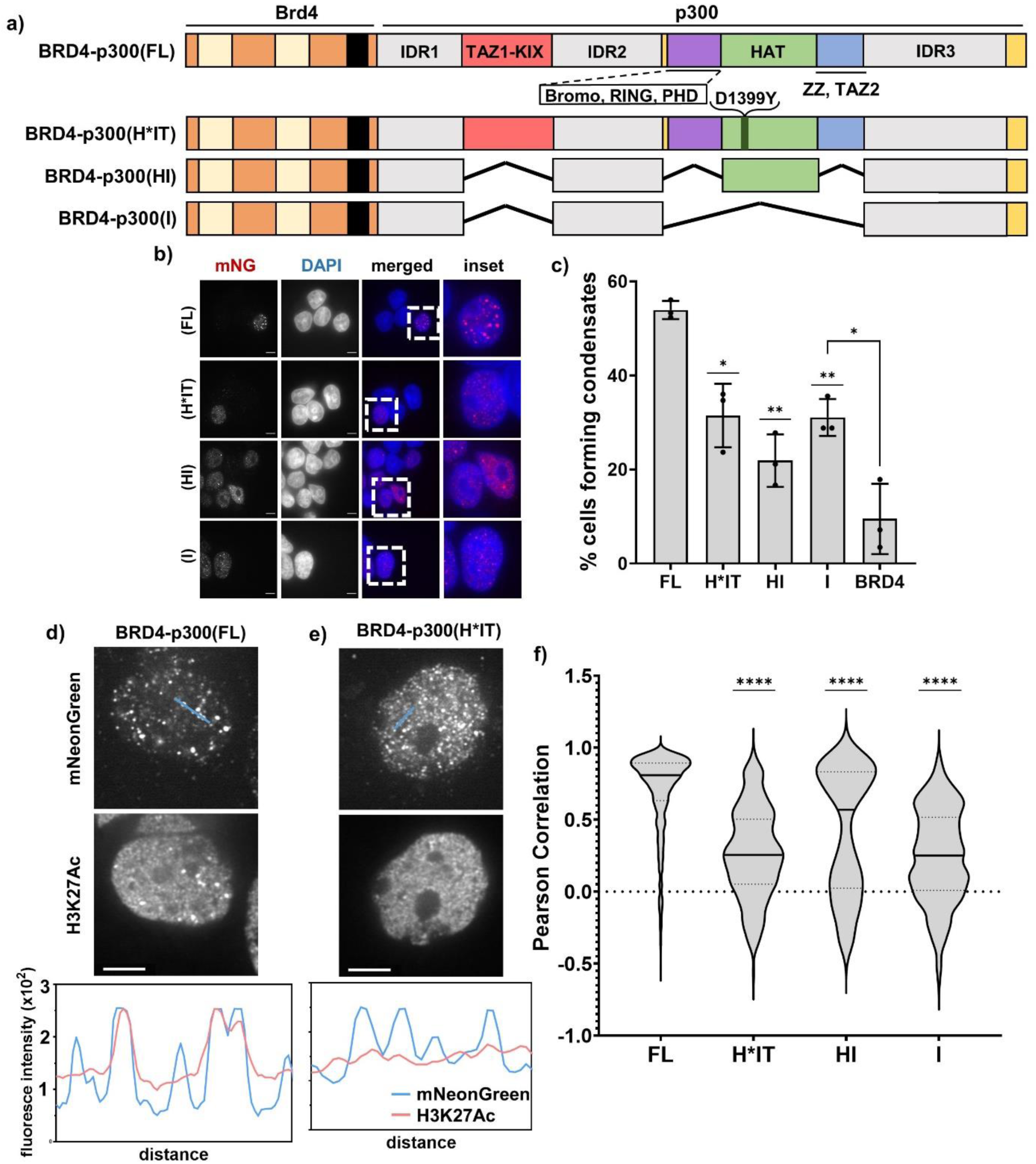
IDRs, HAT and TF binding domains of p300 collectively contribute to condensate formation. a) Schematic of BRD4-p300 – based constructs, including FL, H*IT, HI and I. b) Micrographs of all the BRD4-p300 constructs from a); scale bars = 10μm. c) Quantification of the micrographs represented in b). d) Colocalization of BRD4-p300(FL), as visualized by α-mNeonGreen staining, with α-H3K27Ac staining, shown in micrographs and via line profiles. e) Colocalization of BRD4-p300(H*IT), as visualized by α-mNeonGreen staining, with α-H3K27Ac staining, shown in micrographs and via line profiles. f) Quantification of correlation between α-mNeonGreen and α-H3K27Ac condensates, as shown via Pearson Correlation.

Having established that p300 acetyltransferase activity contributes to BRD4-NUT condensate formation, we next sought to examine the localization of acetylation in cells expressing different BRD4-p300 constructs. There is modest overlap between immunostaining with α-mNeonGreen and α-H3K27Ac antibodies in cells expressing BRD4-p300 (Fig. 7d). Conversely, expression of a HAT-deficient mutant [BRD4-p300(H*IT)] results in a lower level of colocalization, as shown via line profiles (Fig. 7e). When we measured the colocalization between mNeonGreen and H3K27Ac signals in stable cell lines expressing all BRD4-p300 fusion mutants, we discovered that the cells expressing BRD4-p300 and BRD4-p300(HI) have higher levels of colocalization than either of the HAT-deficient mutants [BRD4-p300(H*IT) and BRD4-p300(I)] (Fig. 7f). These results suggest that acetylation is likely concentrated in the condensates due to HAT activity. Together, these data support the idea that the IDRs, HAT and TF-binding all contribute to BRD4-p300 condensate formation, but HAT activity is needed for maximal acetylation within the condensates. By extrapolation, the data suggest that these same dependencies might be found in condensate formation by the complex of p300 with BRD4-NUT.

### P300 acetyltransferase activity is important for transcriptional changes in cells

Finally, we sought to understand if, and how, changes in condensate formation and acetylation are reflected in transcription. We first analyzed the trends in differential gene expression (RNAseq) in stable cell lines expressing the BRD4-p300 mutant fusion proteins, using 293TRex-FlpIn cells as a control. Removal of HAT activity results mostly in gene downregulation [Fig. 8a, ∼74% of all differentially expressed genes for BRD4-p300(H*IT) and ∼60% for BRD4-p300(I)], whereas presence of HAT [BRD4-p300(HI)] results in the majority of genes being upregulated (Fig. 8a, ∼54%). This finding is consistent with the idea that hyperacetylation results in transcriptional activation. We also analyzed the expression of two NC signature genes (*SOX2* and *TP63*) and discovered that they are both significantly upregulated upon expression of all constructs with active HAT [BRD4-p300 and BRD4-p300(HI)] (Fig. 8b). In contrast, in stable cell lines with HAT-deficient BRD4-p300 mutations [BRD4-p300(H*IT) and BRD4-p300(I)], *SOX2* is downregulated and *TP63* is only slightly upregulated, but this latter effect is not statistically significant (Fig. 8b). The third signature gene of NC, *MYC*, was not observed to be upregulated in our stable cell lines, as previously described in other non-NC cell lines expressing BRD4-NUT [8, 21]. This suggests that p300 HAT activity contributes substantially to transcriptional activation by the fusion proteins. Lastly, we applied principal component analysis (PCA) to the RNAseq data to compare transcription changes upon expression of all the constructs (Fig. 8c). Expression profiles for cells expressing BRD4-p300(H*IT) and BRD4-p300(I) cluster most proximally to those of control (239TRex-FlpIn) cells, especially in the PC1 dimension, which carries the highest variance in the data (Fig. 8c). In contrast, cells expressing BRD4-p300 or BRD4-p300(HI) are located farther from control cells in both dimensions and relatively close to each other (Fig. 8c). Thus, the presence of an active HAT domain is sufficient to make the transcriptional profiles resemble the cells expressing full-length BRD4-p300. The presence or absence of TF-binding domains does not have a major effect on transcription, as signified by the close proximity of data points from BRD4-p300(H*IT) and BRD4-p300(I) – expressing cells (Fig. 8c)

**Fig. 8:**
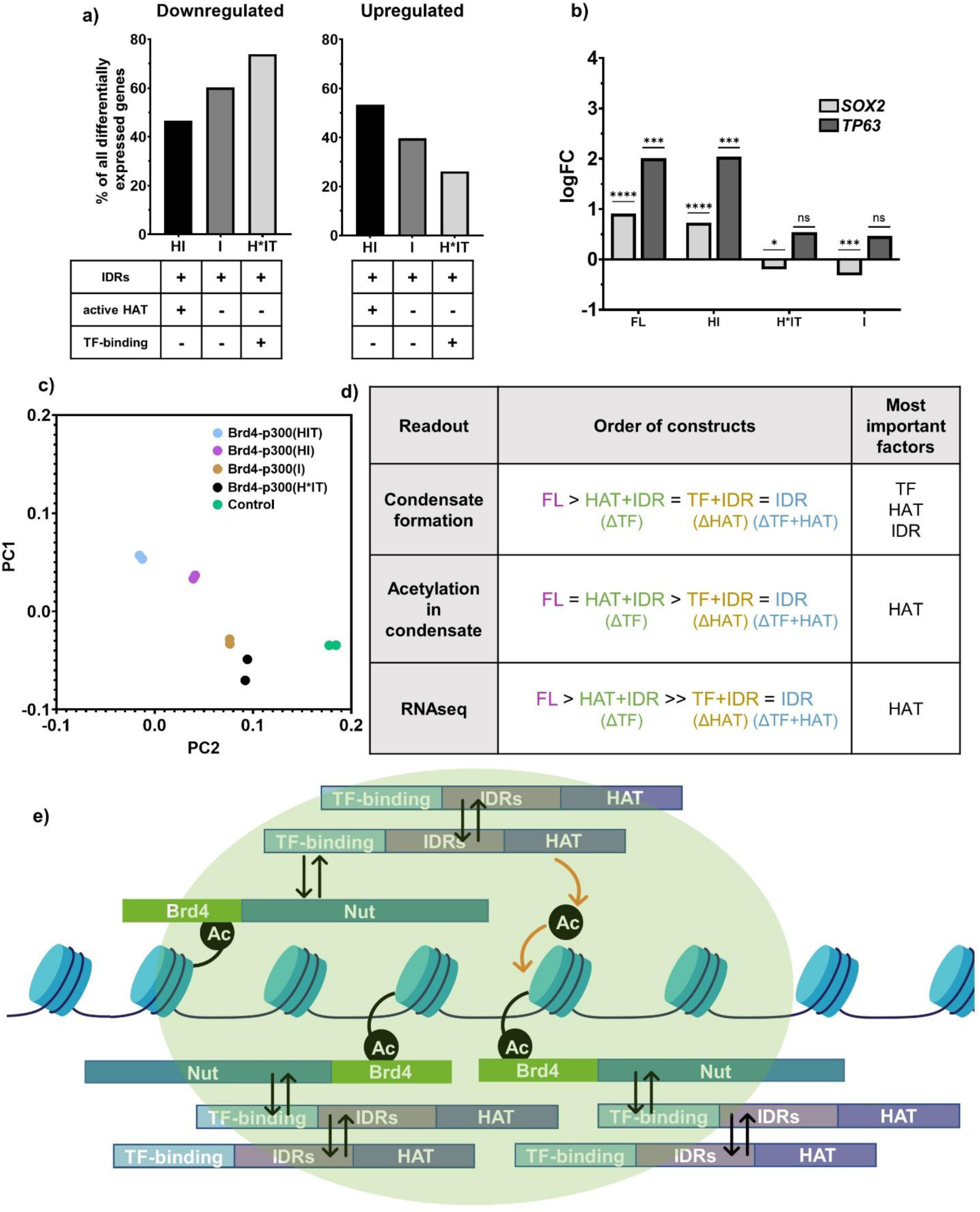
Acetylation activity of p300 is necessary for transcriptional changes observed upon expression of BRD4-p300(FL). a) Percent of genes affected upon expression of different BRD4-p300 constructs. b) Graph showing logFC for *SOX2* and *TP63* genes upon expression of different BRD4-p300 constructs. c) PCA plot, summarizing RNAseq data for cells expressing all constructs. Cell lines excluded in this plot: BRD4-NUT(FL) and BRD4-NUT(MIN) are included in the full PCA in Fig.S4b. d) Summary of molecular parts of p300 required for condensate formation, condensate-localized acetylation and transcriptional changes in cells. e) Final model of condensate formation.

Collectively, these data show that while all three molecular features (IDRs, HAT activity and TF-binding) collectively contribute to condensate formation, HAT activity alone is most important for transcriptional changes. The contribution of each of these molecular elements is thus distinguishable in condensate formation and gene regulation for the BRD4-p300 fusion, and likely for the BRD4-NUT:p300 complex as well.

## Discussion

We studied the connection between BRD4-NUT condensate formation and transcriptional regulation in cells. We found that p300 recruitment by BRD4-NUT is necessary and sufficient for condensate formation. By fusing p300 directly to BRD4, as a mimic of the BRD4-NUT:p300 complex, we found that multiple molecular features of p300 appear to collectively contribute to condensation (Fig. 8d). In contrast, the acetylation activity of the p300 HAT domain is most important among p300 elements for transcriptional changes (Fig. 8d). Thus, the protein regions responsible for condensate formation and transcription by the BRD4-NUT:p300 complex overlap but are not identical.

To study condensate formation and gene expression, we developed a series of inducible stable cell lines. As detailed in Supplementary Materials and Methods section, we carefully controlled protein expression levels, ensuring that our engineered system mimics the expression of BRD4-NUT in the Nut Carcinoma cell line, HCC2429, as closely as possible. As in previous analyses of fusion oncoprotein condensates [1, 2], we also focused our analyses on large condensates to increase our ability to distinguish effects of the different proteins investigated.

We found that the p300-interaction motif of NUT (MIN), in fusion with BRD4, is necessary and sufficient to form condensates to a similar degree as BRD4-NUT and drive similar changes in transcription. Given the importance of this interaction to condensate formation, we examined BRD4-p300 fusions as mimics of the complex, as a more readily manipulated and controlled system. We recognize, however, that elements of this system, including the lack of potential dissociation of p300 from NUT and the expression level of p300 as a fusion, may cause deviations from behavior of the natural BRD4-NUT fusion. With this caveat, further dissection of BRD4-p300 revealed that multiple molecular features in p300 (HAT activity, IDRs, and TF-binding domains) collectively contribute to biomolecular condensation. Interestingly, while both TF-binding domains and HAT contribute to condensate formation (BRD4-p300(H*IT) and BRD4-p300(HI) form condensates in fewer cells than BRD4-p300, when both are missing, the effect is not additive (BRD4-p300(I) forms condensates in a similar fraction of cells as BRD4-p300(H*IT) and BRD4-p300(HI)) (Fig. 7c). This indicates that both TF-binding and HAT activity are required for condensate formation. Thus, TF-binding domains along with HAT might be activating a positive feedback loop. In this model, p300 binds directly to BRD4-NUT through the TF-binding domains and in turn, causes hyperacetylation of nearby histone tails through its HAT domain. Hyperacetylation would then result in increased recruitment of additional BRD4-NUT molecules, through interaction with BRD4 bromodomains, thus closing the positive feedback loop (Fig. 8e). This part of our model is in agreement with a previously postulated general model of BRD4-NUT condensate formation [8]. A novel feature of our model is that p300 IDRs also act in an independent, self-association – based mechanism, to enhance condensate formation (Fig. 8e). This aspect is similar to previous models for other fusion oncoproteins, where a phase separation-inducing element is joined to a functional element, causing aberrant condensate formation and functional activation [1, 4, 43, 44]. This potential involvement of two independent types of interaction, engaging three classes of molecular features of p300 would explain why neither deletion of IDRs or TF-binding domains nor inactivation of HAT results in a complete disruption of condensate formation. Such a mechanism would make condensate formation more robust than with either element alone.

The fact that the MIN fragment of NUT responsible for p300 recruitment is necessary and sufficient for condensate formation indicates that p300 is likely to play important functional roles in NC cells. Indeed, the transcriptional profiles of cells expressing BRD4-NUT(MIN) and BRD4-p300 are similar. However, unlike in the case of condensate formation, where IDRs, HAT and TF-binding all collectively contribute, here we determined that only the HAT activity of p300 strongly influences transcriptional output. We conclude that condensate formation and transcriptional changes upon expression of BRD4-NUT or BRD4-p300 might be decoupled to some extent. Importantly, these results demonstrate that it is not condensate formation, per se, that drives transcriptional activity, since p300 mutants lacking an active HAT domain [BRD4-p300(I) and BRD4-p300(H*IT)] form condensates to some extent but produce fewer and different transcriptional changes. In contrast, a p300 mutant with an active HAT domain (BRD4-p300(HI)) forms condensates to a similar extent as HAT-less mutants and regulates transcription similarly to BRD4-p300. Thus, our molecular dissections suggest that the features that contribute to condensate formation need not necessarily be the same features that enable functional changes, such as transcriptional regulation. However, the biologically-relevant molecules in Nut Carcinoma, the BRD4-NUT fusion protein and its interaction partner, full-length p300, collectively stimulate condensate formation and regulate transcription. Together, our data support a model in which the recruitment of active p300 into BRD4-NUT condensates rewires transcription to drive the gene expression program in Nut Carcinoma; however, the specific molecular features responsible for condensate formation and transcription are distinguishable.

## Acknowledgements

We thank Cheng-Ming Chiang for his generous gift of the BRD4-N antibody. ChIPseq and RNAseq sequencing and analyses were performed by the Next Generation Sequencing Core at McDermott Center at UT Southwestern. We acknowledge Anthony Vega for training and assistance with quantitative image analysis, Benjamin Sabari, Christy Fornero and Heankel Lyons for training on ChIPseq sample preparation, and Bryan Gibson and Lynda Doolittle for assistance initiating the project. Research was supported by the Howard Hughes Medical Institute (to M.K.R.), grants from the Welch Foundation (I-1544 to M.K.R.) and the National Institute Of General Medical Sciences of the National Institutes of Health (R35GM141736 to M.K.R.).

## Author contributions

M.K. and M.K.R. designed research; M.K. performed research; M.K., A.K., C.X., and M.K.R. analyzed data; and M.K., S.L.C., and M.K.R. wrote the paper.

## Materials and methods

### Cell Culture

HCC2429 cells were a kind gift from the Hamon Center for Therapeutic Oncology Research at UT Southwestern Medical Center. The cells were grown in RPMI 1640 medium (Thermo Fisher Scientific, #11875119) with addition of 10% heat-inactivated Fetal Bovine Serum (Thermo Fisher Scientific, #10-438-026) and 1% penicillin – streptomycin (Thermo Fisher Scientific, #15140-122). The cell line was validated via immunofluorescence microscopy and western blotting for BRD4-NUT expression. HCC2429 cells were expanded during an early passage after receiving the cell line and multiple aliquots were frozen for future use. The cell line was regularly tested negative for mycoplasma, using the MycoAlert™ Detection Kit (Lonza, #LT07-418) and MycoAlert™ Control Set (Lonza, #LT07-518).

293TRex-FlpIn cells were purchased from Thermo Fisher Scientific (#R78007). The cells were grown in DMEM medium (Thermo Fisher Scientific, #11965-118) with addition of 10% heat-inactivated Fetal Bovine Serum (Thermo Fisher Scientific, #10-438-026), 1% penicillin – streptomycin (Thermo Fisher Scientific, #15140-122), 15µg/mL blasticidin (Thermo Fisher Scientific, #A1113903) and 100µg/mL zeocin (Thermo Fisher Scientific, #R250-01). The cell line was regularly tested negative for mycoplasma, as described above.

293TRex-FlpIn cell lines inducibly expressing different constructs were developed based on the original 293TRex-FlpIn cell line, according to the manufacturer’s instructions. The list of constructs successfully introduced into the cell line are: mNeonGreen-BRD4(short), “BRD4”, mNeonGreen-NUT, “NUT”, mNeonGreen-BRD4-NUT, “BRD4-NUT”, mNeonGreen-BRD4-NUT(355-505), “BRD4-NUT(MIN)”, mNeonGreen-BRD4-NUT(Δ355-505), “BRD4-NUT(ΔMIN)”, mNeonGreen-BRD4-p300, “BRD4-p300”, mNeonGreen-BRD4-p300(D1399Y), “BRD4-p300(H*IT)”, mNeonGreen-BRD4-p300(ΔTF-binding), “BRD4-p300(HI)”, and mNeonGreen-BRD4-p300(IDR-only), “BRD4-p300(I)”. We were not able to develop a stable cell line inducibly expressing mNeonGreen-BRD4-p300(ΔIDR) construct, we think due to high toxicity of the construct. Experiments that present data with use of mNeonGreen-BRD4-p300(ΔIDR) were performed using transient transfections. All cell lines were maintained in DMEM medium (Thermo Fisher Scientific, #11965-118) with addition of 10% heat-inactivated Fetal Bovine Serum (Thermo Fisher Scientific, #10-438-026), 1% penicillin – streptomycin (Thermo Fisher Scientific, #15140-122), 15µg/mL blasticidin (Thermo Fisher Scientific, #A1113903) and 100µg/mL hygromycin B (Thermo Fisher Scientific, #10687010). All the cell lines were expanded in an early passage after developing and frozen in multiple aliquots for future use. Each cell line was tested for protein expression after induction, via immunofluorescence microscopy and western blotting. All cell lines regularly tested negative for mycoplasma, as described above.

### Immunofluorescence

Immunofluorescence was performed with HCC2429 cells, 293TRex-FlpIn cells and all newly developed cell lines described above, with Hoechst 33342 nuclear counterstaining. The primary antibodies used were: NUT monoclonal rabbit antibody (Cell Signaling Technology, #3625S), mNeonGreen monoclonal mouse antibody (Chromotek, #32f6-100), p300 monoclonal mouse antibody (Santa Cruz Biotechnology, #sc-56455), BRD4 monoclonal rabbit antibody (Abcam, #ab128874), BRD4 monoclonal mouse antibody (Abcam, #ab244221) and H3K27Ac monoclonal rabbit antibody (Abcam, #ab4729). Secondary antibodies were: Alexa Fluor 568 – conjugated goat anti-mouse secondary antibody (Thermo Fisher Scientific, #A-11004) and Alexa Fluor 647 – conjugated goat anti-rabbit secondary antibody (Thermo Fisher Scientific, #A21245).

24-well glass bottom microscopy plates (Cellvis, #P24-1.5H-N) were treated with 5µg/mL poly-D-lysine (Sigma-Aldrich, #P7405-5MG) for 1h at room temperature, washed with 1×PBS (Thermo Fisher Scientific, #10010049) and dried for 2h at room temperature in the tissue culture hood.

Cells were seeded 1 day before induction, assuring optimal density for imaging. After induction with doxycycline for 2h, followed by a 4h washout (Sigma-Aldrich, #D9891-5G), cells were washed with 1mL PBS per well and fixed with 0.5mL 4% paraformaldehyde (Thermo Fisher Scientific, #RT15710) for 20 minutes at room temperature. Next, the cells were washed 3 times with 1mL PBS per well for 3 minutes. Cells were then permeabilized with 1mL per well of 0.5% Triton-X-100 (Thermo Fisher Scientific, #J66624AP) in PBS for 10 minutes at room temperature and then washed 3 times with 1mL PBS per well for 3 minutes. Next, the cells were blocked in the blocking buffer – 5% BSA (Thermo Fisher Scientific, #BP9704100) in PBST (PBS with 0.1% Tween-20 (Sigma-Aldrich, #P7949-500ML)) for 1 hour at room temperature. Primary antibodies were diluted in the blocking buffer as follows: NUT monoclonal rabbit antibody – 1:500, mNeonGreen monoclonal mouse antibody – 1:750, p300 monoclonal mouse antibody – 1:500, BRD4 monoclonal rabbit antibody – 1:500, H3K27Ac monoclonal rabbit antibody – 1:2000. The cells were incubated with 300µL of the primary antibody dilution overnight at 4°C. The cells were then washed 3 times, for 5 minutes each time, with 1mL of PBST per well. Appropriate secondary antibodies were diluted 1:1000 in the blocking buffer. The cells were incubated with 300µL of the secondary antibody dilution for 2 hours at room temperature, during which the plate was covered with aluminum foil to protect the samples from light. Maintaining the protection from light, the cells were next washed 3 times for 5 minutes with 1mL of PBST per well. Next, the cells were incubated with 500µL of 1:2000 dilution of Hoechst 33342 (Thermo Fisher Scientific, #H3570) in PBS for 10 minutes at room temperature. Finally, the cells were washed 3 times for 3 minutes with 1mL of PBS and then, 1mL of fresh PBS was used as a mounting medium for each well. Finally, a 1:100 dilution of the secondary antibodies in PBS were prepared and 500µL of such dilutions were placed in separate empty wells, for flat-field correction imaging. One well was always prepared with PBS only, for background imaging. Plates prepared in this way were kept at 4°C and covered with aluminum foil until performing microscopy.

### Confocal Microscopy

All samples were imaged on a Nikon Eclipse Ti microscope base equipped with a Yokogawa CSUX1 spinning disk confocal scanner unit, using 100× / 1.49 NA oil objective and Andor EM-CCD camera. Images were acquired using MetaMorph software. A single Z-slice in the center of cell nucleus was acquired per image, with an average of 100-200 images acquired per sample. The lasers used were: 405nm, 488nm, 561nm and 637nm.

### Quantitative Microscopy – Analyses

For microscopy data analysis, nuclear segmentation was achieved using Cellpose software with the preset nuclear diameter of 80. Downstream microscopy analyses were performed via CellProfiler software version 4.2.1 for Windows, using the following sequence of modules:

- *CorrectIlluminationApply* – to subtract background (empty image) from the empty image with fluorophore. To obtain the empty images, we used an empty well in microscopy plate with 1×PBS, as imaging medium. To obtain the empty image with fluorophore, we made serial dilutions of Alexa Fluor – conjugated secondary antibodies used in immunostaining of cells and tested them all. Based on empirical analysis, I decided to use a 1:100 dilution of the antibodies as the fluorescent background.
- *CorrectIlluminationCalculate* – to calculate the illumination function across the corrected illumination image after background subtraction. Here, we determined a mathematical description of illumination pattern across a micrograph and created an image that is representative of the overall illumination pattern. This calculation was performed on the background-subtracted empty image, created using the module described above. The illumination function is later used to correct for a potential uneven illumination in images.
- *RescaleIntensity* – to divide the illumination function image by its maximum (now the maximum intensity of the illumination function is 1).
- *CorrectIlluminationApply* – to subtract background (empty image) from the image with cells, in the same way that it was done in the first background subtraction step described above.
- *CorrectIlluminationApply* – to divide the background-subtracted image of cells by the illumination function image after corrections, determined in the steps described above. This produces a flat-field corrected image of cells.
- *IdentifyPrimaryObjects* – we uploaded the DAPI stain-based segmented images produced through Cellpose into CellProfiler and masked them. We used a typical nucleus diameter of 80-300 pixels, to make sure all nuclei are included, and removed any nuclei that are touching the image borders. The best thresholding method for nuclei identification in the masks from Cellpose was empirically determined to be Otsu.
- *EnhanceOrSuppressFeatures* – to enhance the fluorescence intensity of pixels within condensates relative to the rest of the image, resulting in improvement of subsequent identification of condensates.
- *MaskImage* – to mask the nuclei in the image with cells using the segmented images produced through Cellpose.
- *IdentifyPrimaryObjects* – to identify the condensates – round I: using a first, more lenient method of thresholding with a typical condensate diameter of 6-40 pixels, adaptive thresholding with the Robust Background thresholding method and size of adaptive window of 50 pixels, determined empirically. Adaptive thresholding methods calculate a different threshold for each pixel, thus adapting to potential differences in fore-and background fluorescence intensities in the analyzed image. This step allowed us to identify all large condensates, but often does not account for their correct shapes, merging low intensities surrounding condensates or combining some of the large condensates with neighboring small condensates.
- *MaskImage* – to mask out the condensates identified using the module above.
- *IdentifyPrimaryObjects* – to identify the condensates – round II: using a second, more stringent method of thresholding: typical condensate diameter: 8-40 pixels, global thresholding performed with the global Otsu thresholding method with a threshold correction factor: 0.5. Global thresholding methods calculate a single threshold value for the input image. Then, they use this determined value to classify pixels that have intensities higher than the threshold as foreground and the ones with lower intensity as background. Since the condensates have already been masked, following round I of identification, we empirically determined that the global thresholding method works well to account for the large condensates’ shapes and correctly splits neighboring condensates that might otherwise have been merged in the first round of identification.
- *MeasureObjectIntensity* – to measure the fluorescence intensity across the nuclei as well as within the identified condensates. The quantitative fluorescence intensity information obtained is reported as average within the nucleus, average among condensates within a given cell and average within a given condensate. Through this module, we can subsequently apply the fluorescence intensity cutoffs in our analyses, as described in Fig.S1 and S2.
- *MeasureObjectSizeShape* – to measure the condensate size and shape: through this module, we determined the eccentricity of condensates, which is a numerical descriptor of roundness. This is an important measure of morphology differences among different constructs.
- *RelateObjects* – to establish a relationship between nuclei and condensates: here, the algorithm records which condensates are found in which nucleus.
- *MeasureObjectOverlap* – to quantify the overlap between different channels, e.g., in the case of co-immunostaining with two separate antibodies, such as α-BRD4 and α-p300 or α-NUT and α-histone H3K27Ac etc.
- *MeasureObjectColocalization* – to quantify the overlap between different features found in all channels, e.g., condensates found in a co-immunostaining experiment. One of the methods utilized here is Pearson Correlation, which can be calculated based on the fluorescence intensity overlap between condensates identified in two different channels. One channel is used as a ground truth and fluorescence intensities from condensates identified in another channel are compared to it.

Each module was optimized, based on a set of representative micrographs. All modules were run in the order listed.

### Statistical analyses

Welch’s t-test was used for pairwise comparisons in figures: 1d, 3c, 4a, 5c, 6d, 7c and S6b. Kruskall-Wallis nonparametric test with Dunn’s multiple comparisons of ranks between preselected pairs was performed in figures: 4b and 5d. Kolmogorov-Smirnov nonparametric test, comparing cumulative distributions was performed in figures: 7f and S6d.

### Cell cross-linking for ChIP

Cells were grown in 15cm dishes to confluency and 2-3 plates were used for each experiment. After doxycycline treatments, cells were washed once with 20mL of PBS. Next, the cells were treated with 1% methanol-free formaldehyde (Thermo Scientific, #28908) diluted in PBS – 10mL per plate for 8 minutes, with gentle agitation. Formaldehyde was then quenched with 500µL of 2.5M glycine (to a final concentration of 125mM) for another 8 minutes, with gentle agitation. After quenching, cells were scraped off the plates, transferred to 50mL conical tubes and centrifuged at 500×g at 4°C for 5 minutes. The supernatant was removed, and the cell pellet was washed with 5mL of cold PBS per plate. Following the wash, cells were again centrifuged at 500×g at 4°C for 5 minutes and then, the supernatant was removed. Pellets were finally snap-frozen in liquid nitrogen and stored at –80°C.

### Chromatin extraction and shearing

Cell pellets were thawed and resuspended in LB1 buffer (50mM Hepes-KOH pH 7.9, 140mM NaCl, 1mM EDTA, 10% glycerol, 0.5% NP40, 1% TritonX-100 and 1× Complete, EDTA-free protease inhibitor cocktail, Sigma Aldrich, #11873580001) to obtain ∼1×10^7^ cells/mL suspension and they were then incubated on a rotator at 4°C for 20 minutes, for lysis. Lysis efficiency of at least 60% was determined using trypan blue. Next, the suspension was centrifuged at 1,350×g at 4°C for 5 minutes. The supernatant was removed, and the pellet was resuspended in LB2 buffer (10mM Tris pH 8.0, 200mM NaCl, 1mM EDTA, 0.5mM EGTA and 1× Complete, EDTA-free protease inhibitor cocktail) to again obtain ∼1×10^7^ cells/mL suspension and they were then incubated on a rotator at 4°C for 5 minutes. Next, the suspension was centrifuged at 1,350×g at 4°C for 5 minutes. The supernatant was removed, and the pellet was resuspended in LB3 buffer (10mM Tris pH 8.0, 100mM NaCl, 1mM EDTA, 0.5mM EGTA, 0.1% sodium deoxycholate, 0.5% SDS, 1% TritonX-100 and 1× Complete, EDTA-free protease inhibitor cocktail) to obtain ∼1×10^7^ cells/mL suspension, which was then passed through a 27G needle 3 times, to completely homogenize the pellet. The suspension was transferred to a Covaris millitube with AFA fiber (Fisher Scientific, # NC0597431) and sonicated using Covaris M220 sonicator (average incident power 7.5 Watts, peak incident power 75 Watts, duty factor 10%, cycles: 200 count, duration: 18 minutes, minimum temperature: 5°C, temperature setpoint: 7°C, maximum temperature: 9°C). After sonication, samples were centrifuged at 15,000×g at 4°C for 10 minutes, soluble supernatant was transferred to a new tube, snap-frozen in liquid nitrogen and stored at –80°C. 10µL of each sample were incubated overnight at 65°C and treated with Proteinase K and RNAse A and DNA was purified to test the sonication efficiency (details described below, in “Chromatin Immunoprecipitation” part).

### Chromatin immunoprecipitation

Protein G-conjugated Dynabeads (Thermo Fisher Scientific, #10004D) (75uL of suspension per IP) were washed with 1mL of blocking buffer (0.5% BSA in PBS) 3 times and collected on a magnet stand each time. Beads were resuspended in 500µL blocking buffer and mixed with 7.5µg antibody (NUT monoclonal rabbit antibody (Cell Signaling Technology, #3625S), or H3K27Ac monoclonal rabbit antibody (Abcam, #ab4729)) and incubated on a rotator at 4°C overnight. Next day, the beads were washed 3 times with the same blocking buffer to remove unbound antibody and then they were resuspended in 75µL of the blocking buffer. The resuspended beads bound to antibody were mixed with chromatin extract and mixed on a rotator at 4°C overnight. After that, the beads were washed 3 times in 1mL washing buffer 1 (50mM Hepes pH 7.0, 100mM LiCl, 1mM EDTA, 1% NP40, 0.7% sodium deoxycholate and 1× Complete, EDTA-free protease inhibitor cocktail). For the third wash, beads were incubated on a rotator at 4°C for 10 minutes. The washes were repeated with washing buffer 2 (20mM Tris pH 8.0, 350mM NaCl, 1% TritonX-100, 0.1% SDS, 2mM EDTA). Next, the beads were washed with 1mL TE buffer with 50mM NaCl and centrifuged shortly at 300×g and then placed on the magnet stand, to remove residual buffer. Then, 200µL of the elution buffer was added to the beads (50mM Tris pH 8.0, 10mM EDTA and 1% SDS) and the beads were incubated at 65°C for 30 min. with gentle agitation. The beads were next centrifuged at 300×g and placed on the magnet stand. The 200µL elution was then transferred to a new tube. The elution was incubated at 65°C overnight to reverse crosslinks. Next, samples were treated with RNase A (Thermo Scientific, #EN0531) for 1h at 37°C and then with Proteinase K (Fisher Scientific, #25-530-049) for 2h at 55°C. DNA was then purified from these samples using the Qiagen PCR purification kit (Qiagen, #28106).

### ChIP data processing and representation

Libraries were sequenced on Illumina NextSeq 500 at Next Generation Sequencing Core, Eugene McDermott Center for Human Growth and Development).

For alignment, single-end reads with a read length of 100 bp were generated for each library. FASTQ files were checked for quality using fastqc (v0.11.5, https://www.bioinformatics.babraham.ac.uk/projects/fastqc/) and fastq_screen (v0.11.4, https://www.bioinformatics.babraham.ac.uk/projects/fastq_screen/). Low-quality reads and adapter were removed using fastq-mcf (v1.05, http://expressionanalysis.github.io/ea-utils/). The reads from FASTQ files were aligned to the human genome (hg19) using bowtie2 (v2.3.3.1) [45]. Picard-tool’s (v2.10.10 https://broadinstitute.github.io/picard/) MarkDuplicates module was then used to remove duplicated alignments.

For peak calling, annotation and motif analysis, the duplicate removed alignment files were used to call peaks using MACS2 (v2.1.0) [46], with a q-value threshold of 0.05 and using DNA input as background controls. The fragment size of each library was used to extend reads at their 3′ ends to a fixed length with “–extsize” parameter in MACS2. The peak files from the peak calls were annotated using annotatePeaks module in HOMER [47].

ChIPseq tracks shown were visualized using Integrative Genomics Viewer [48, 49]. Venn diagrams were generated using BioVenn [50, 51]. The representation factor is the number of overlapping genes divided by the expected number of overlapping genes, when randomly drawn from two independent groups.

### RNA sequencing sample preparation

Cells were grown in 10cm plates to confluency. After treatment with doxycycline, cells were lysed and RNA was purified using the Qiagen RNeasy kit (Qiagen, # NC9677589). Purified RNA was stored at –20°C until use.

### RNA sequencing data processing and representation

Samples were sequenced on the Illumina NextSeq 500 with read configuration as 75 bp, single end reads. (This part should be include in the RNA library preparation section towards the end).

The Fastq files were subjected to quality check using fastqc (v0.11.5, http://www.bioinformatics.babraham.ac.uk/projects/fastqc) and fastq_screen (v0.11.4, http://www.bioinformatics.babraham.ac.uk/projects/fastq_screen). The reads from FASTQ files were aligned to the human genome (hg19) using STAR (v2.5.3a) [52].

For differential gene expression, read counts were generated using featureCounts [53] from the Rsubread package (v1.4.6). Differential expression analysis was performed using edgeR [54]. Statistical cutoffs of *p-value* < 0.05 and log2FC > 2 were used to identify statistically significant differentially expressed genes.

Venn diagrams were generated using BioVenn [50]. Heatmaps for RNAseq-ChIPseq integration were generated using Heatmapper [55].

### Western Blotting

For Western Blotting, cells were grown to confluency in 6-well plates or 10cm dishes induced with doxycycline. Cells were then washed once with PBS and lysed with RIPA buffer, supplemented with 400mM NaCl for 30 minutes. Clarified cell lysate was tested via BCA assay (Thermo Fisher Scientific, #PI23227) to quantify the protein concentration. After adjusting the concentrations of samples, 15µL of samples were mixed with 15µL of 2× SDS buffer and heated on a 100°C heating block for 5-10 minutes. Next, the samples were loaded on a 10% SDS-PAGE gel made in-lab or on a 4-15% pre-cast TGX gel (Bio-Rad, #4568086), along with the molecular marker. Gels were run for 40 minutes – 1h at 200V. Transfer onto a PVDF membrane (Sigma Aldrich, #: IPVH00010) was performed wet, using a transfer buffer with addition of 0.1% SDS. After transfer, membrane was blocked for 1h at room temperature using blocking buffer (5% BSA in TBST). Next the membrane was incubated with the appropriate primary antibody in blocking buffer overnight at 4°C. The following primary antibodies were used for Western Blotting: rabbit anti-BRD4(N) antibody (generous gift from the Chiang lab at UT Southwestern), rabbit anti-NUT (C52B1) antibody (Cell Signaling Technology, #3625S), mouse anti-p300 antibody (Millipore Sigma, #05-257), mouse anti-GAPDH (Thermo Fisher Scientific, #MA515738). Next day the membrane was washed 3 times, for 5 minutes each with TBST. Then the membrane was incubated with the appropriate secondary antibody in the blocking buffer at room temperature for 2h. The following secondary antibodies were used for Western Blotting: mouse anti-rabbit HRP antibody (Santa Cruz Biotechnology, #sc-2357) and goat anti-mouse HRP antibody (Thermo Fisher Scientific, #31430). Next, the membrane was washed 2 times 10 minutes. in TBST and once for 10 minutes. in TBS. For signal development, the antibody was removed and membrane was incubated for about 3 minutes. in a solution with chemiluminescent substrate (Fisher Scientific, #WBKL S0 100).

### mNeonGreen pulldown

To perform the mNeonGreen pulldowns, the cells were grown on a 10cm dish and treated with doxycycline for induction. Next, the cells were lysed with RIPA buffer supplemented with 400mM NaCl. For the pulldown, we purchased the mNeonGreen-Trap Agarose Kit (Proteinteck, #ntak) and followed the manufacturer’s instructions. After the pulldown, the samples were ran on a gradient, 4-15% gel (Bio-Rad, #4568086). Transfer and Western Blotting was performed as described above, in the “Western Blotting” section.

### Molecular biology and cloning

The following constructs were cloned into the pcDNA5/FRT/TO vector, purchased from Thermo Fisher Scientific (#V652020): mNeonGreen-BRD4(short), named “BRD4”, mNeonGreen-NUT, named “NUT”, mNeonGreen-BRD4-NUT, named “BRD4-NUT”, mNeonGreen-BRD4-NUT(355-505), named “BRD4-NUT(MIN)”, mNeonGreen-BRD4-NUT(Δ355-505), named “BRD4-NUT(ΔMIN)”, mNeonGreen-BRD4-p300, named “BRD4-p300”, mNeonGreen-BRD4-p300(D1399Y), named “BRD4-p300(H*IT)”, mNeonGreen-BRD4-p300(ΔTF-binding), named “BRD4-p300(HI)”, and mNeonGreen-BRD4-p300(IDR-only), named “BRD4-p300(I)” and mNeonGreen-Brd4-p300(ΔIDR). Some of the constructs were initially introduced into pInducer20 vector, and then moved into the pcDNA5/FRT/TO vector. The inserts were amplified using the following primers:

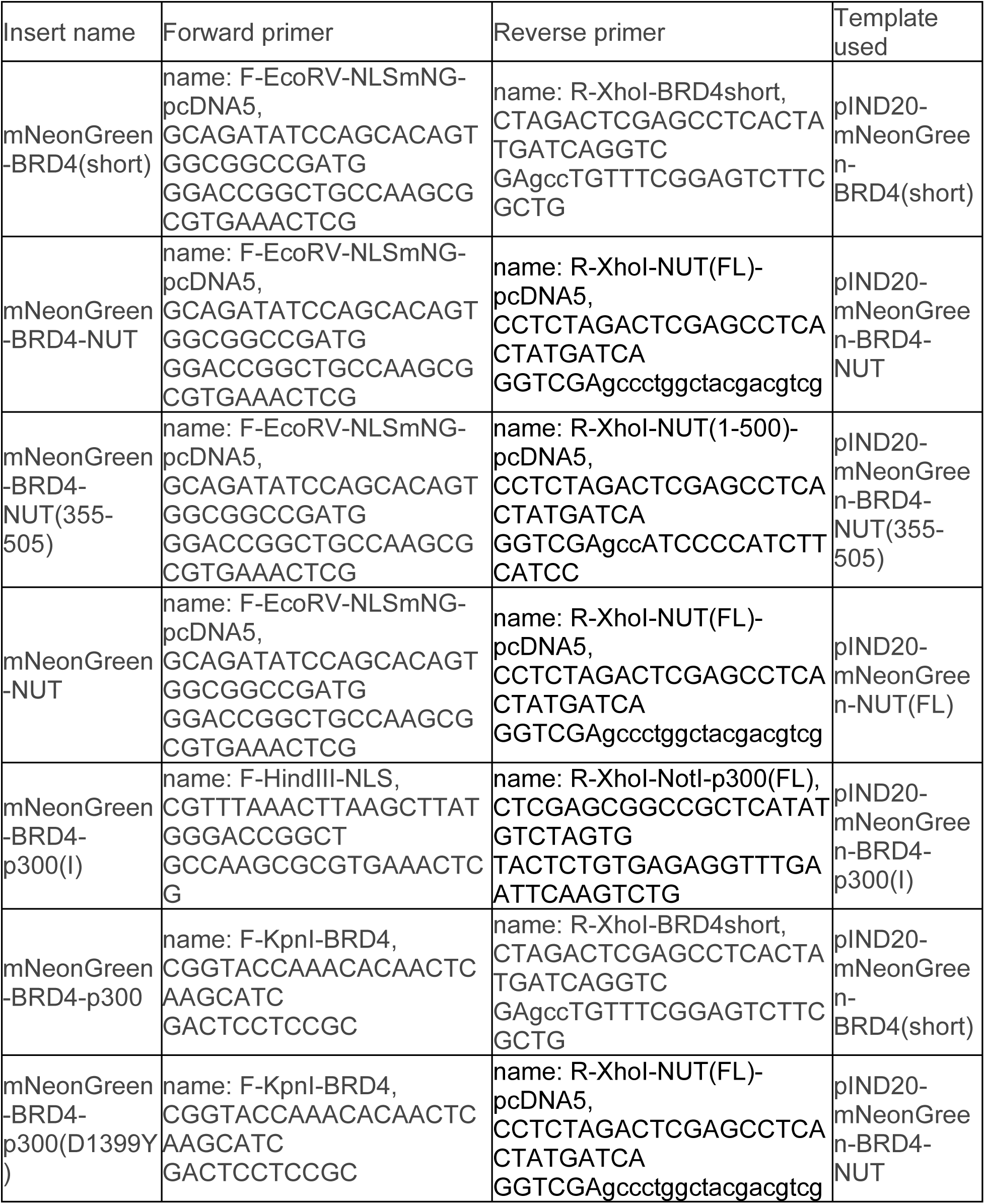
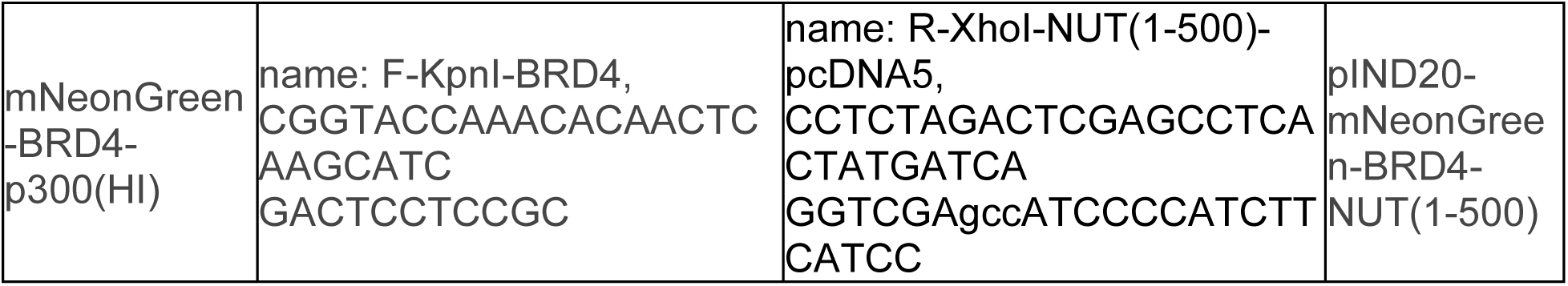

Amplification was performed using Pfu Turbo polymerase (Agilent, #600252), according to the manufacturer’s instructions. After successful amplification, DNA was PCR-purified using Qiagen PCR purification kit (Qiagen, #28106). The amplified fragments and the pcDNA5/FRT/TO vector were then digested with the enzymes listed in the names of primers in the table above. Digestion was performed at 37°C for 1-4h, using Cut Smart buffer from NEB, supplied along with the enzymes. After that time, Quick CIP phosphatase (NEB, # M0525S) was added to the vector and the mix was incubated at 37°C for additional 10-15 minutes. The fragments were then ligated into the vector using T4ligase (Enzymatics, # L6030-LC-L) and transformed into Max Efficiency Stbl2 Competent Cells (Thermo Fisher Scientific, #10268019). The following constructs were obtained using exactly this method:

- mNeonGreen-BRD4(short)
- mNeonGreen-BRD4-NUT
- mNeonGreen-BRD4-NUT(355-505)
- mNeonGreen-NUT
- mNeonGreen-BRD4-p300(I)

For the remaining constructs, cloning was performed in two rounds, because the inserts to be ligated into the vector were larger than the vector itself. First, a 1366bp fragment was ligated into the vector, using the following primers:

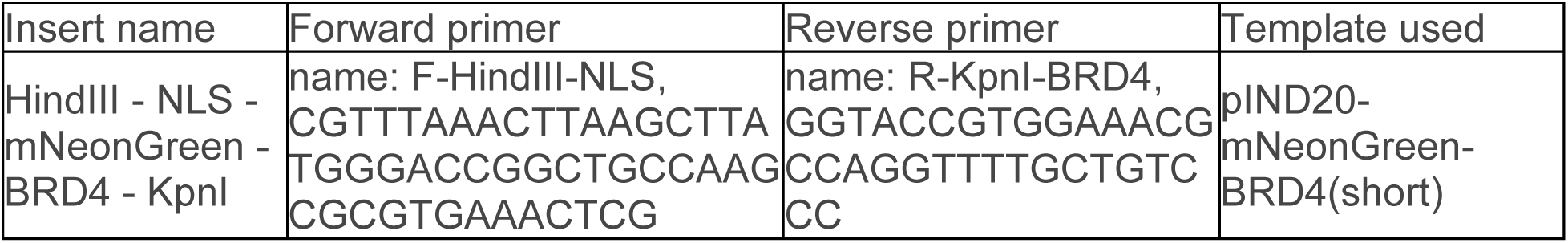

The amplified fragment and the pcDNA5/FRT/TO vector were both digested using HindIII and KpnI enzymes, using the method described above. After obtaining and sequencing the new vector with the 1366bp insert, the remaining inserts were amplified, using the primers listed in the first table:

- mNeonGreen-BRD4-p300
- mNeonGreen-BRD4-p300(H*IT)
- mNeonGreen-BRD4-p300(HI)

The digest of the new vector (with 1366bp fragment insert) and digest of the new inserts was performed using the specified enzymes, in the method described above. The fragments were ligated with T4 ligase and DNA was transformed into the Stbl2 bacteria. After DNA purification all inserts were fully sequenced.

Finally, the mNeonGreen-BRD4-NUT(ΔMIN) construct was obtained using Gibson assembly protocol, with the following primers:

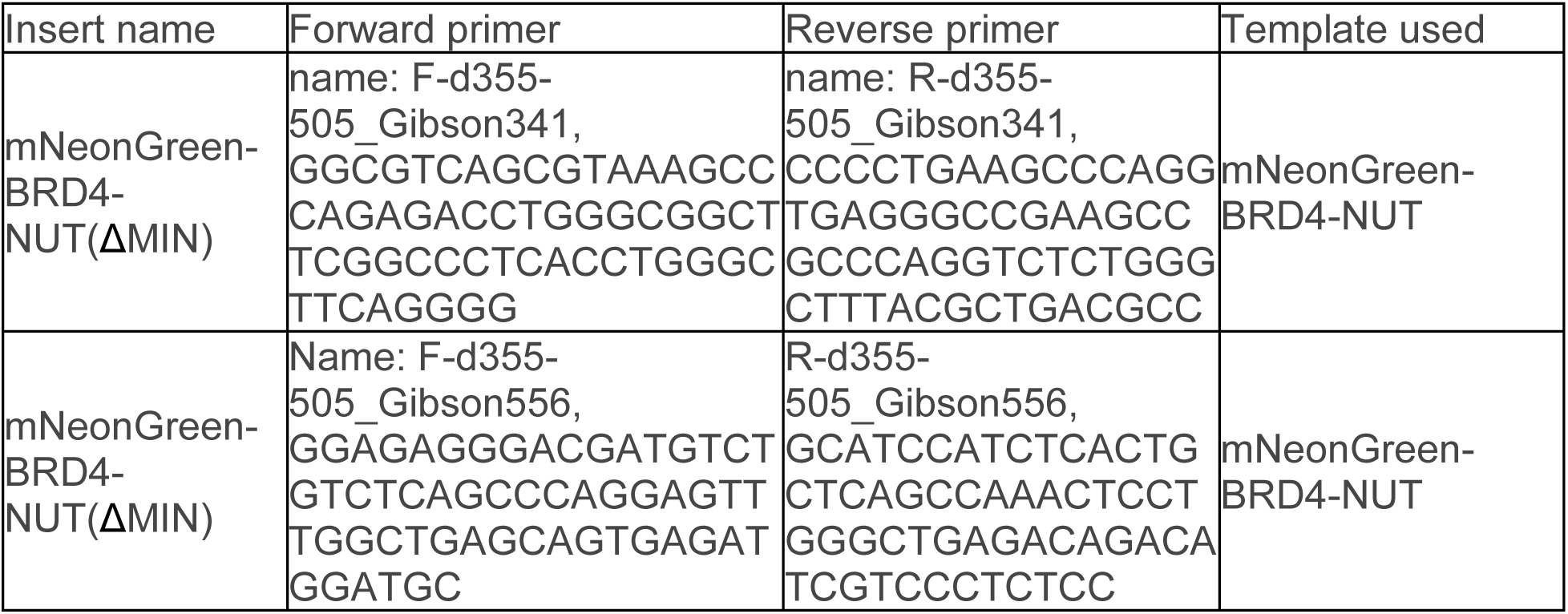

After DNA purification the construct was sequenced.

