## Supplemental Material for "Molecular features driving condensate formation and gene expression by the BRD4-NUT fusion oncoprotein are overlapping but distinct"

**Suppl. Fig. 1: Doxycycline – induced protein expression varies from cell to cell but is very similar between Nut and mNeonGreen antibody staining.**

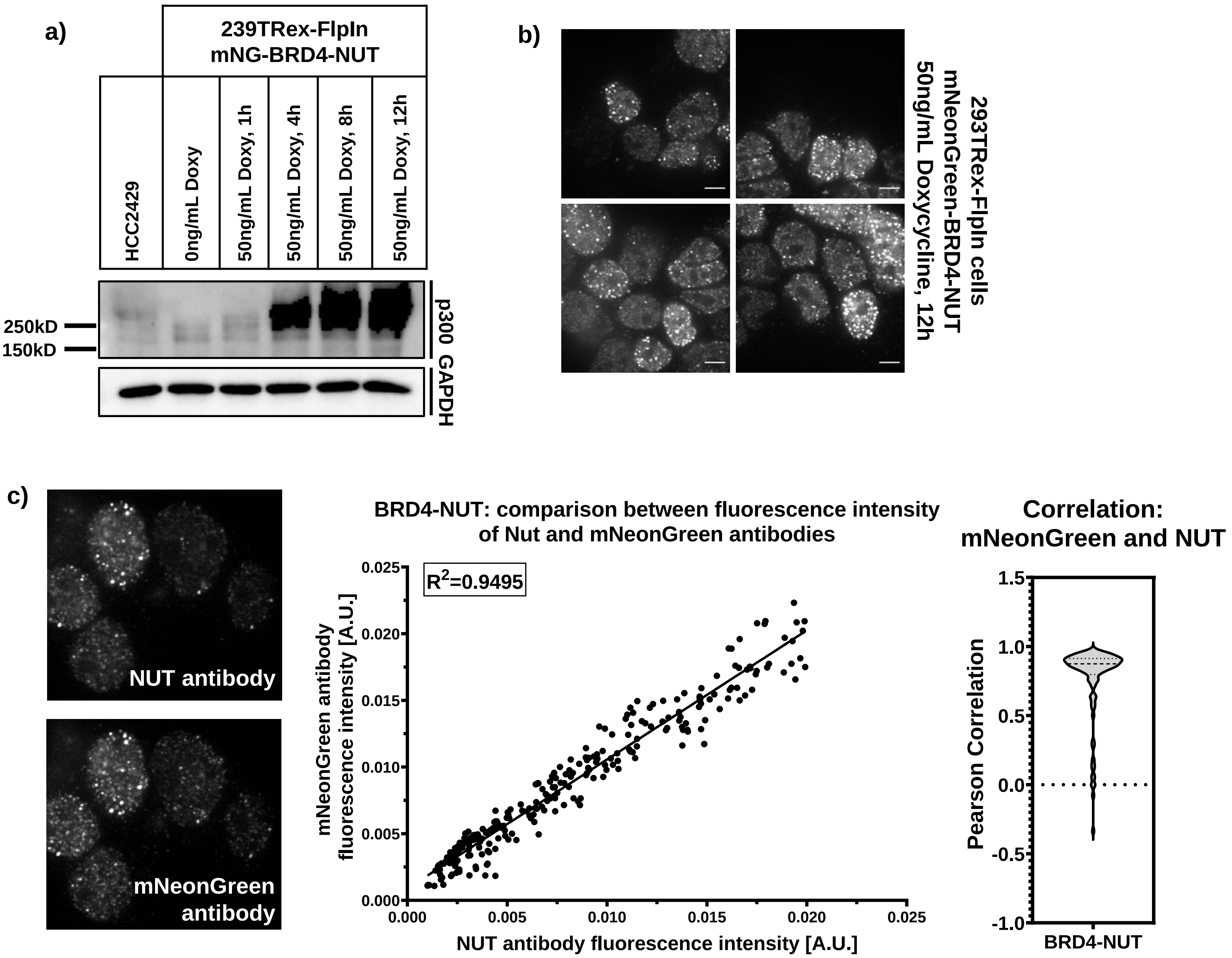

**Suppl. Fig. 1: Doxycycline – induced protein expression varies from cell to cell but is very similar between NUT and mNeonGreen antibody staining.**

a) Western blot showing the difference in average expression level upon 1-12h treatment with 50ng/mL Doxycycline.

b) Example images of cells that have been treated with 100ng/mL Doxycycline for 12 hours – expressing mNG-BRD4-NUT(FL) – notice the big expression differences within each field of view. Scale bar = 10µm

**c)** **Left**: micrographs of representative cells expressing BRD4-NUT(FL) co-stained with mNeonGreen and NUT antibody; staining shows high level of colocalization between the two channels. **Center**: fluorescence intensity of mNeonGreen antibody in relation to fluorescence intensity of NUT antibody; signal intensity highly correlates ($R^2$=0.9495). **Right**: Pearson correlation between the condensates found via immunostaining with mNeongreen antibody or NUT antibody.

Because there is a very high correlation between the two antibodies staining and the resulting fluorescence intensities, both these antibodies are used in quantitative analyses.

**Suppl. Fig. 2: Controlling doxycycline-induced protein expression levels for comparing condensate formation between cells expressing different constructs.**

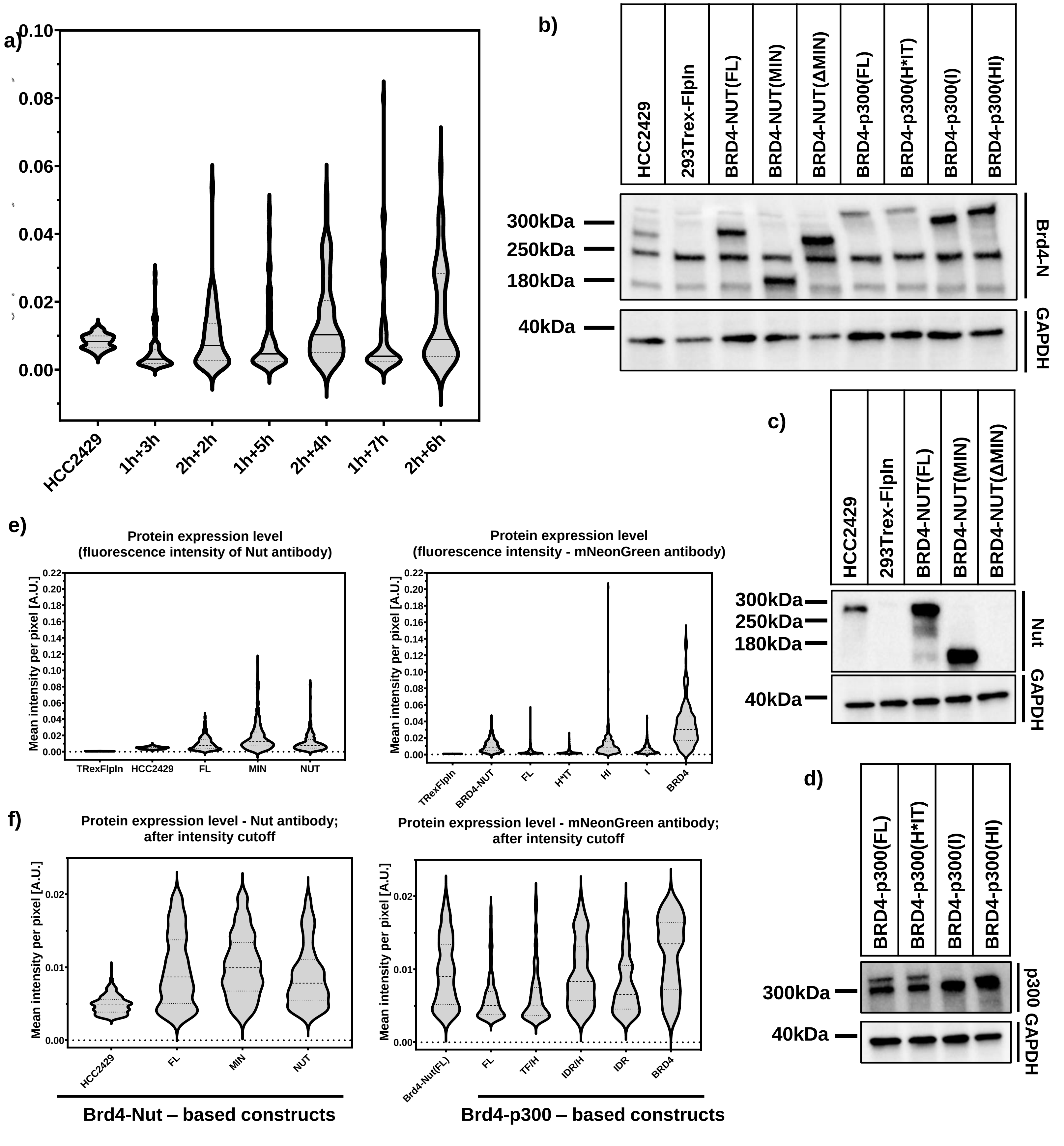

**Suppl. Fig. 2: Controlling doxycycline-induced protein expression levels for comparing condensate formation between cells expressing different constructs.**

a) Protein expression level as measured by average pixel intensity in HCC2429 carcinoma cell line and BRD4-NUT(FL) – expressing stable cell line after 1h treatment with 5ng/mL Doxycycline followed by 3, 5 or 7h washout or after 2h Doxy treatment followed by 2, 4 or 6h washout, as labeled, where e.g. 1h+3h means 1h treatment with Doxy followed by 3h washout. We chose 2h treatment, followed by 4h washout for a good dynamic range and values similar to HCC2429.

b) Western blot showing similarities and differences in the expression levels between cell lines expressing different constructs. All cell lines treated with 5ng/mL Doxycycline for 2h, followed by 4h washout. Antibodies used include: BRD4(N) and GAPDH as a loading control. All fusion protein constructs are expressed as relatively similar levels, with only BRD4-p300(FL) and BRD4-p300(H*IT) expressing slightly lower than others.

c) Western blot showing similarities and differences in the expression levels between cell lines expressing different BRD4-NUT – based constructs. Antibodies used include NUT and GAPDH as a loading control. Note that the epitope for NUT antibody is likely located within the MIN fragment of NUT, as the antibody does not stain the BRD4-NUT(ΔMIN) construct.

d) Western blot showing similarities and differences in the expression levels between cell lines expressing different BRD4-p300 – based constructs. Antibodies used include p300 and GAPDH as a loading control. Note that the size difference between wild-type p300 and either BRD4-p300(I) or BRD4-p300(HI) constructs is too small to see the bands from both.

e) Expression level of fusion proteins in all imaged cell lines, as measured by average fluorescence intensity, using Nut antibody or mNeonGreen antibody; stable cell lines treated with 5ng/mL Doxycycline for 2h, followed by a 4h washout.

f) Expression levels shown upon setting an experimentally established expression cutoff at 0.003-0.02. The cutoff was set to resemble the expression of BRD4-NUT in HCC2429 cell line as well as possible; stable cell lines treated with 5ng/mL Doxycycline for 2h, followed by a 4h washout.

**Suppl. Fig. 3: NUT is a large, mostly disordered protein with a few patches of predicted α-helical structures.**

**a)**

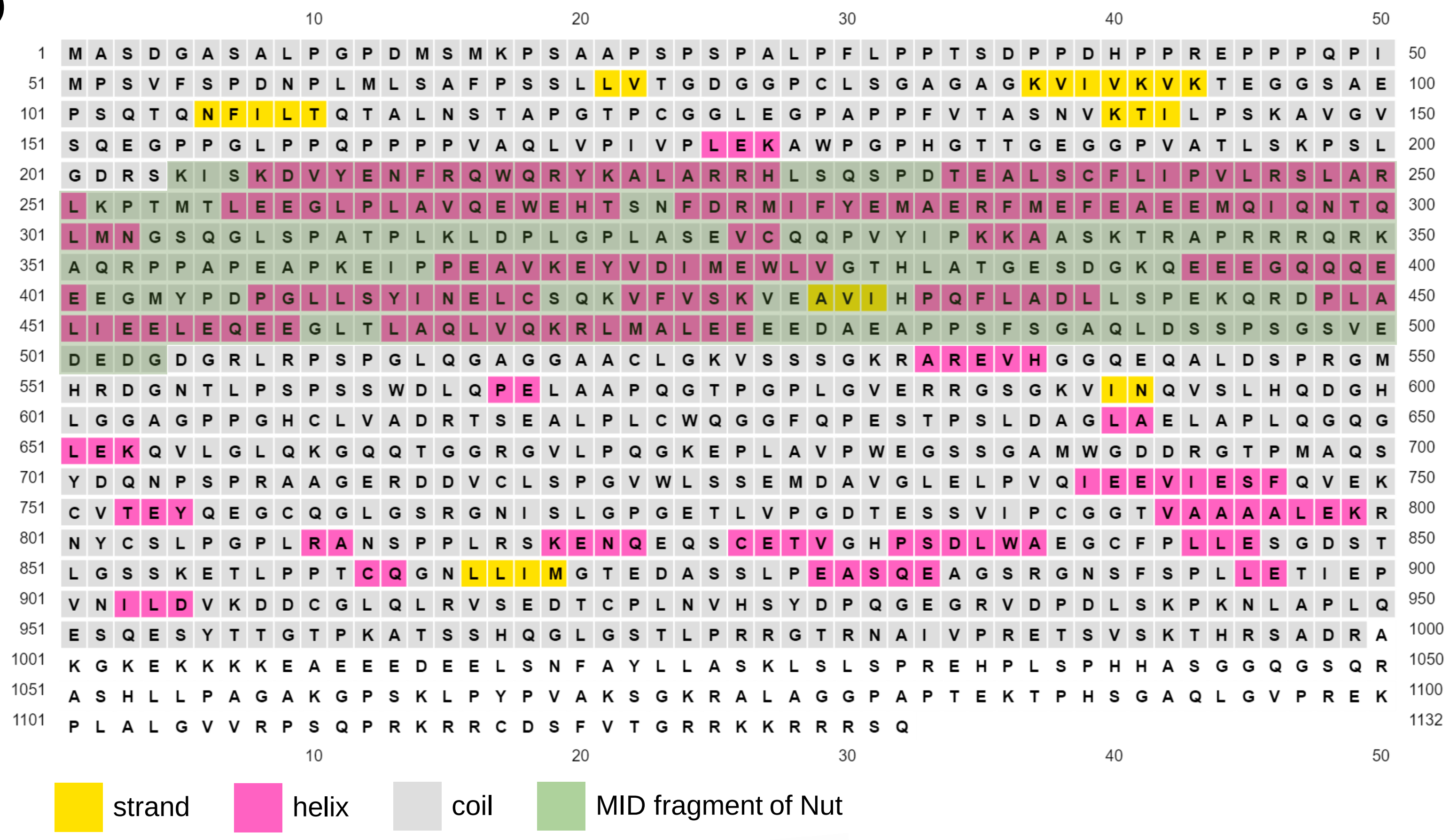


**b)**

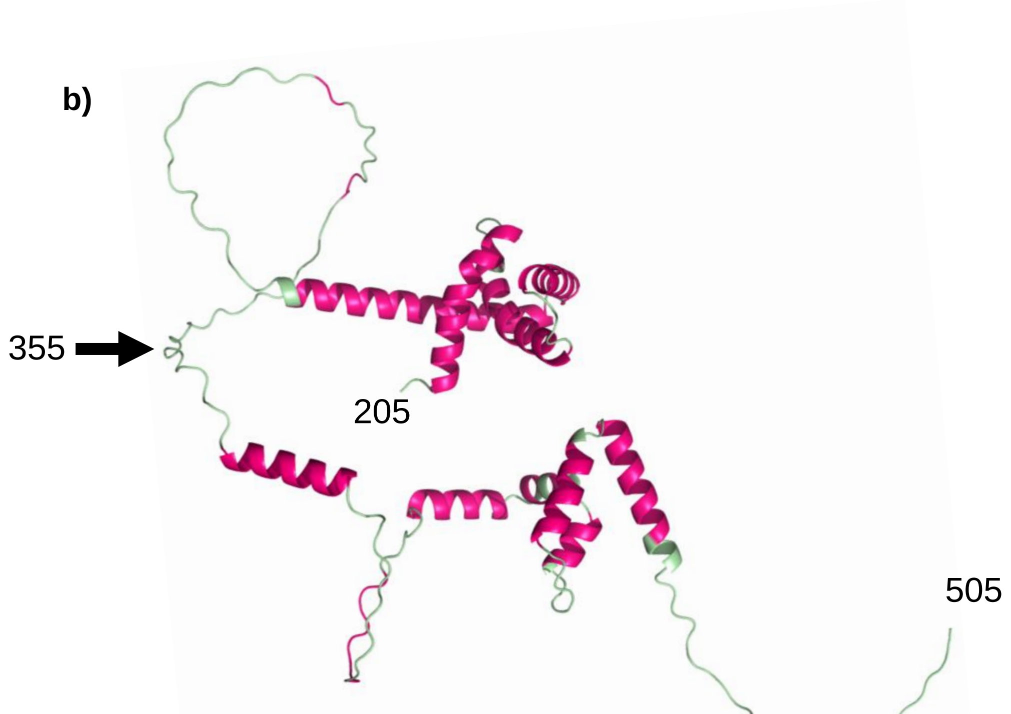


NUT-MID (205-505)

NUT-MIN (355-505)

**Suppl. Fig. 3: NUT is a large, mostly disordered protein with a few patches of predicted α-helical structures.**

a) Psi-pred secondary structure prediction of NUT, with fragments used in MIN and MID – containing constructs highlighted in green.

b) AlphaFold2 secondary structure prediction of NUT(MID), color coded to match the Psi-pred prediction in a).

**Suppl. Fig. 4: Transcription and gene occupancy correlation between different cell lines.**

a)

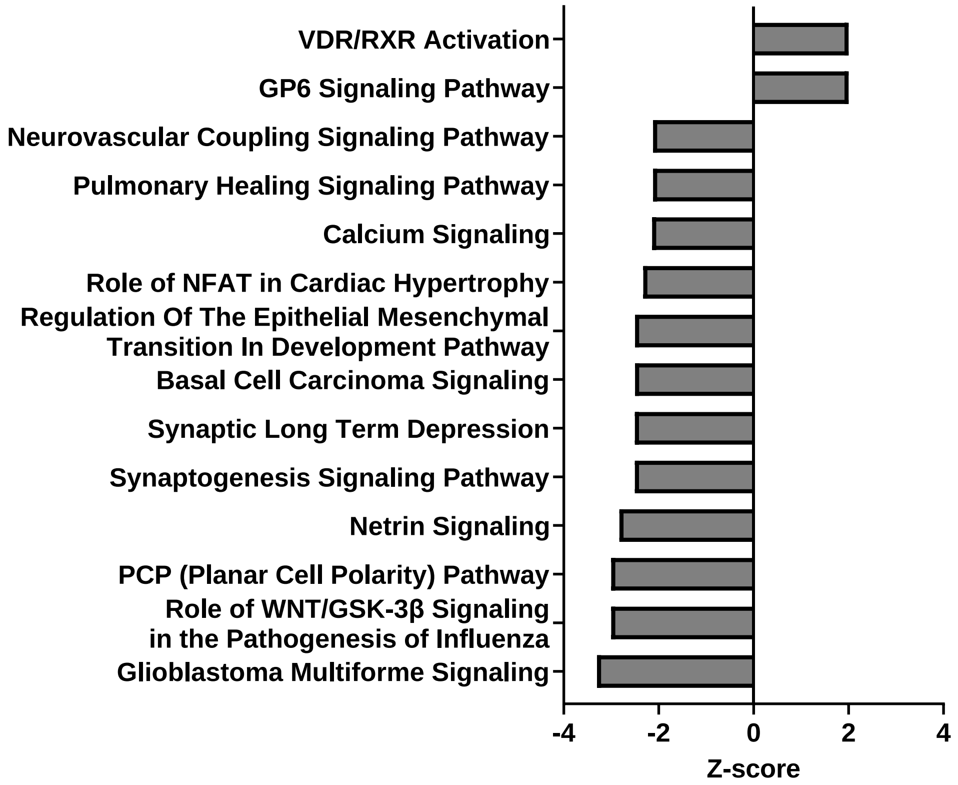


b)

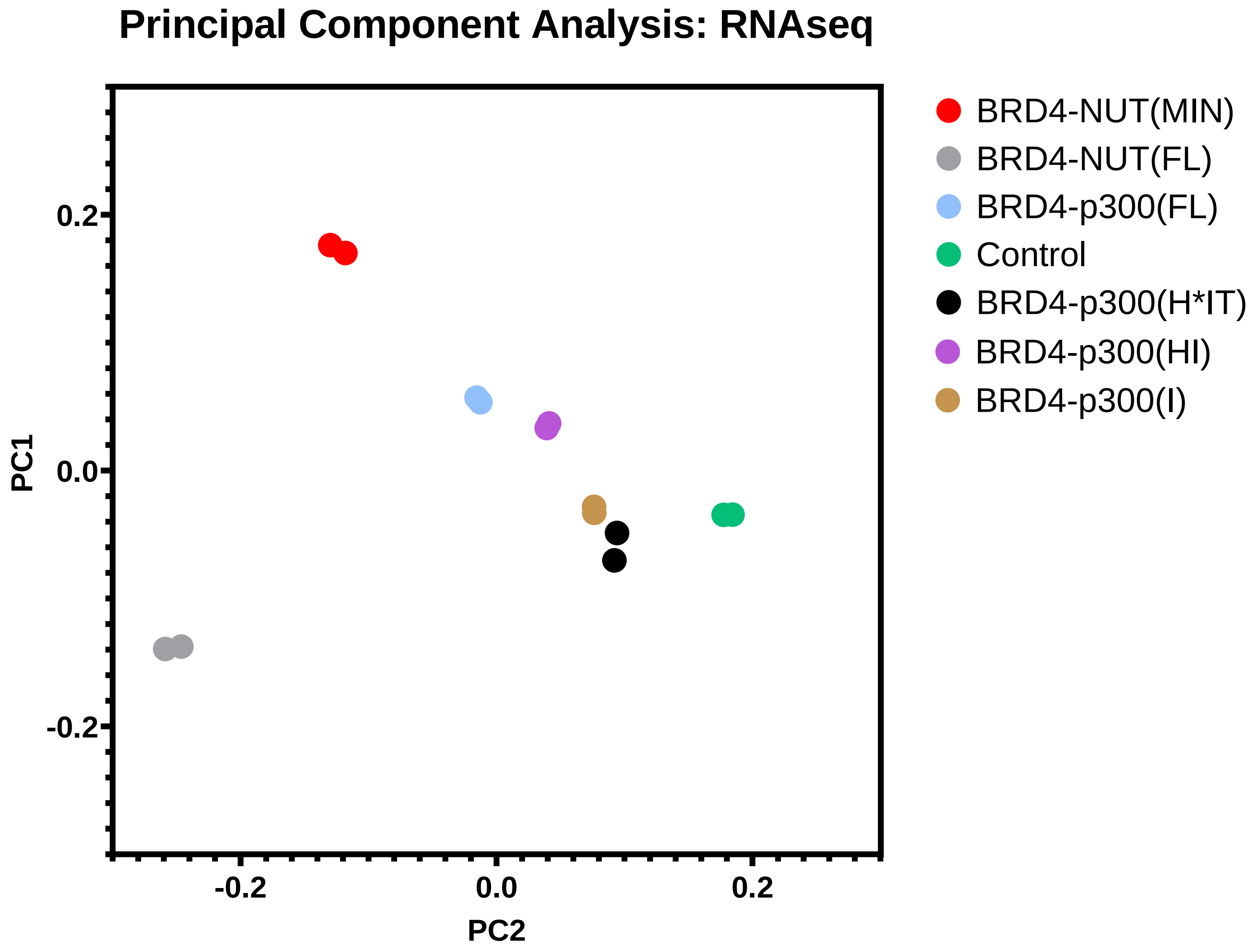

**Suppl. Fig. 4: Transcription and gene occupancy correlation between different cell lines.**

a) Ingenuity Pathway Analysis: most significantly up- and downregulated pathways, based on RNAseq of cells expressing BRD4-NUT(FL), as compared to 293TRex-FlpIn cells not expressing any fusion protein. Z-score is a measure of up- or downregulation. Only pathways with a p value of $< 0.05$ and Z-score $\geq 2$ or $\leq (-2)$ are reported.

b) PCA plot, summarizing RNAseq data for cells expressing all constructs. Subset of this plot, only including Brd4-p300 mutant expressing cell lines is shown in Fig. 8d.

**Suppl. Fig. 5: CIDER and PSPredictor analyses of p300 IDRs propensity to phase separate**

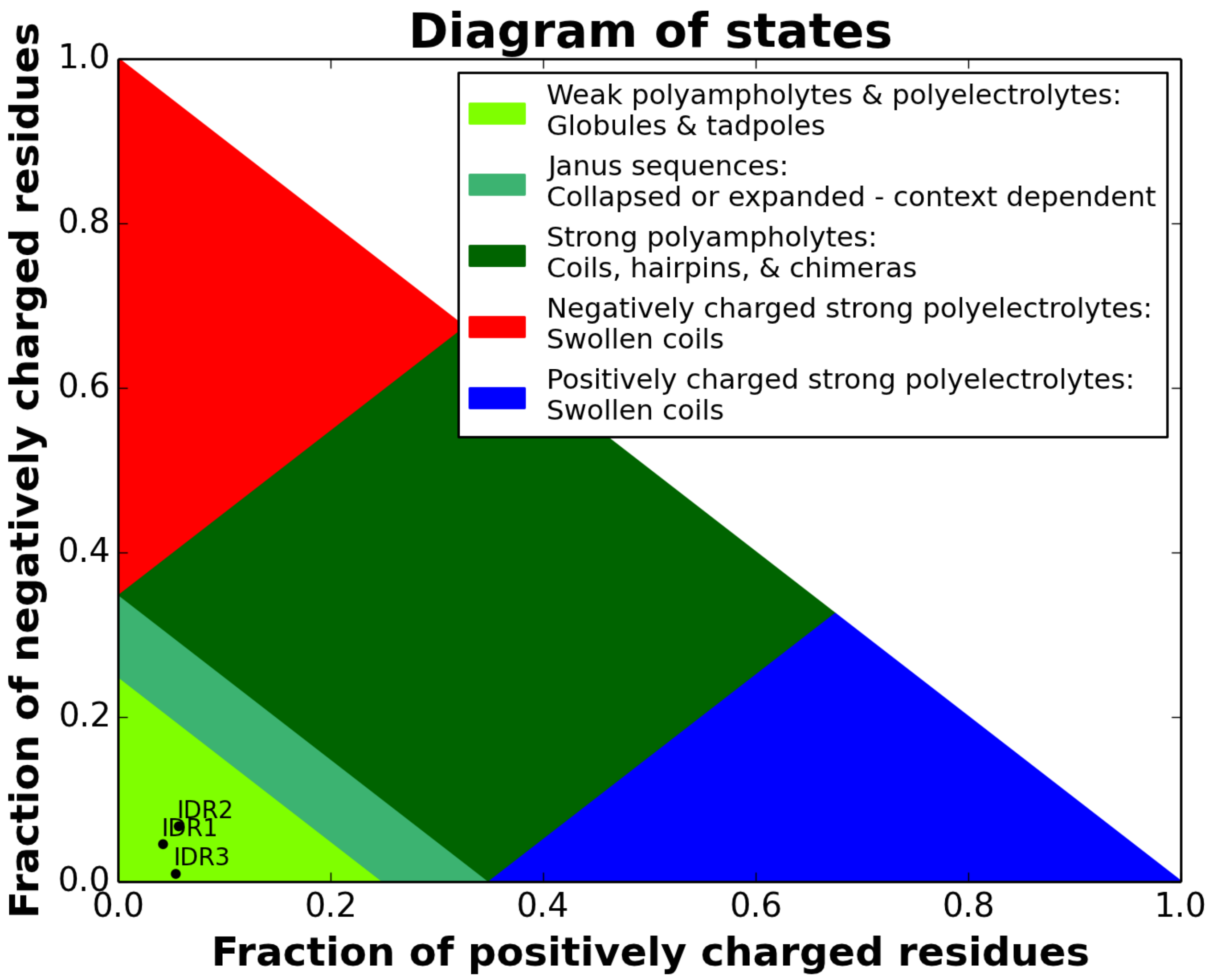


| PSPredictor prediction results | | |
|---|---|---|
| Seq ID | PSP Score | PSP (Yes/No) |
| IDR1 | 0.9573 | Yes |
| IDR2 | 0.9995 | Yes |
| IDR3 | 0.9973 | Yes |

**Suppl. Fig. 5: CIDER and PSPredictor analyses of p300 IDRs propensity to phase separate:**

All three IDRs of p300 classify as “weak polyampholytes and polyelectrolytes” in CIDER analysis and are predicted to be prone to phase separate based on their sequence.

**Suppl. Fig. 6: Applying a stringent expression level cutoff allows to compare condensate formation between stable cell lines and cells expressing constructs transiently.**

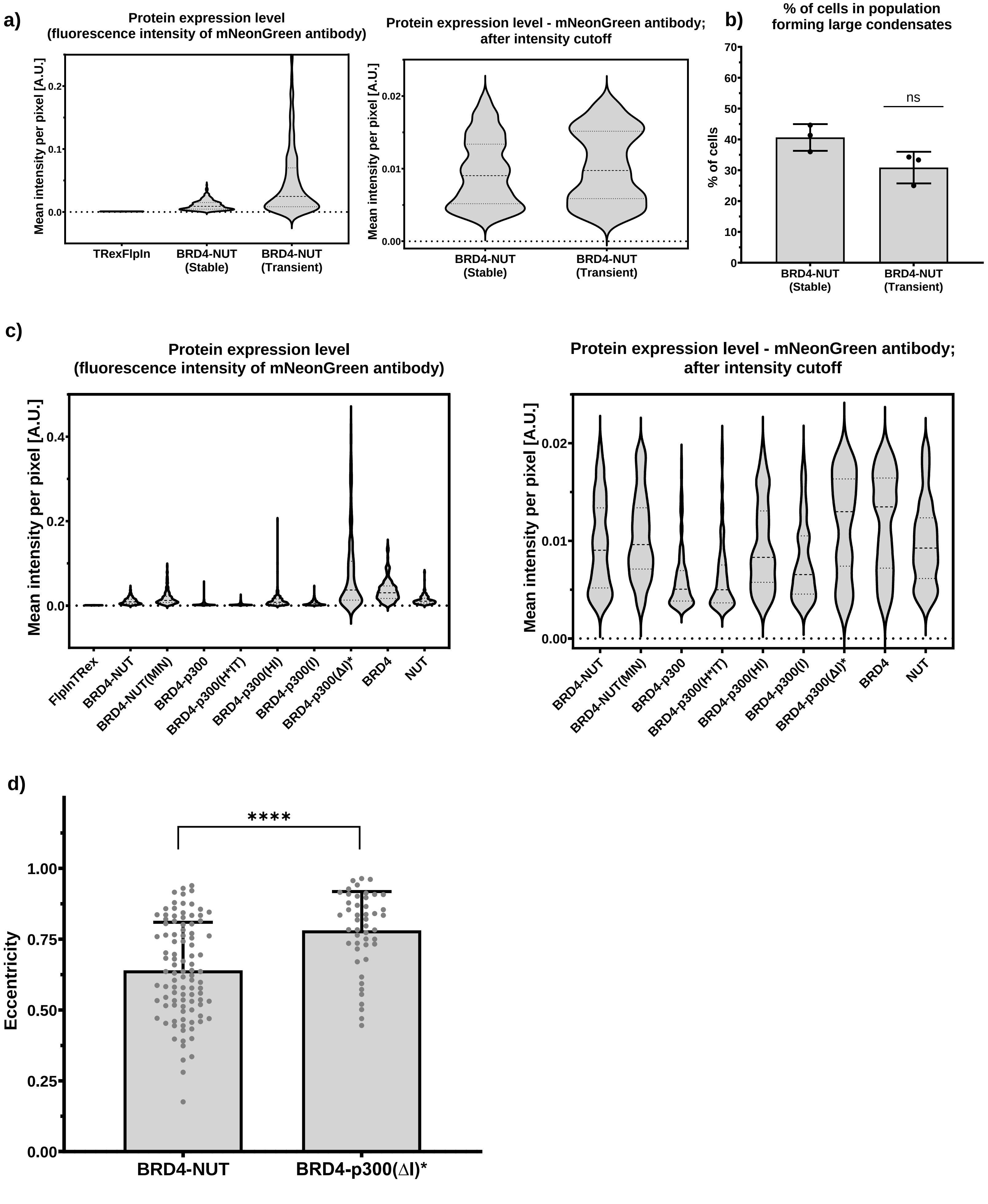

**Suppl. Fig. 6: Applying a stringent expression level cutoff allows to compare condensate formation between stable cell lines and cells expressing constructs transiently.**

a) Quantification of protein expression level as shown by average pixel intensity, to compare the difference between a stable cell line and cells expressing the same construct transiently. Left side graph shows all data and right-side graph shows the intensities recorded after applying a fluorescence intensity cutoff.

b) Percentage of cells forming large condensates: comparison between the stable cell line and transiently transfected cells. The data in transiently transfected cells are shown after applying the expression level cutoff from a).

c) Protein expression level among different stable cell lines, as shown by average pixel intensity, to compare the difference between stable cell lines and the cells expressing Brd4-p300(ΔIDR) transiently. Left side graph shows all data and right-side graph shows the intensities recorded after applying the fluorescence intensity cutoff.

d) Eccentricity of condensates formed by Brd4-Nut(FL) construct and Brd4-p300(ΔIDR) construct. The graph shows that the Brd4-p300(ΔIDR) condensates are less round than the ones formed by Brd4-Nut(FL).

**Suppl. Fig. 7: Analysis of a potential bleed through between laser channels and cross-reactivity between the α-NUT and α-mNeonGreen antibodies**

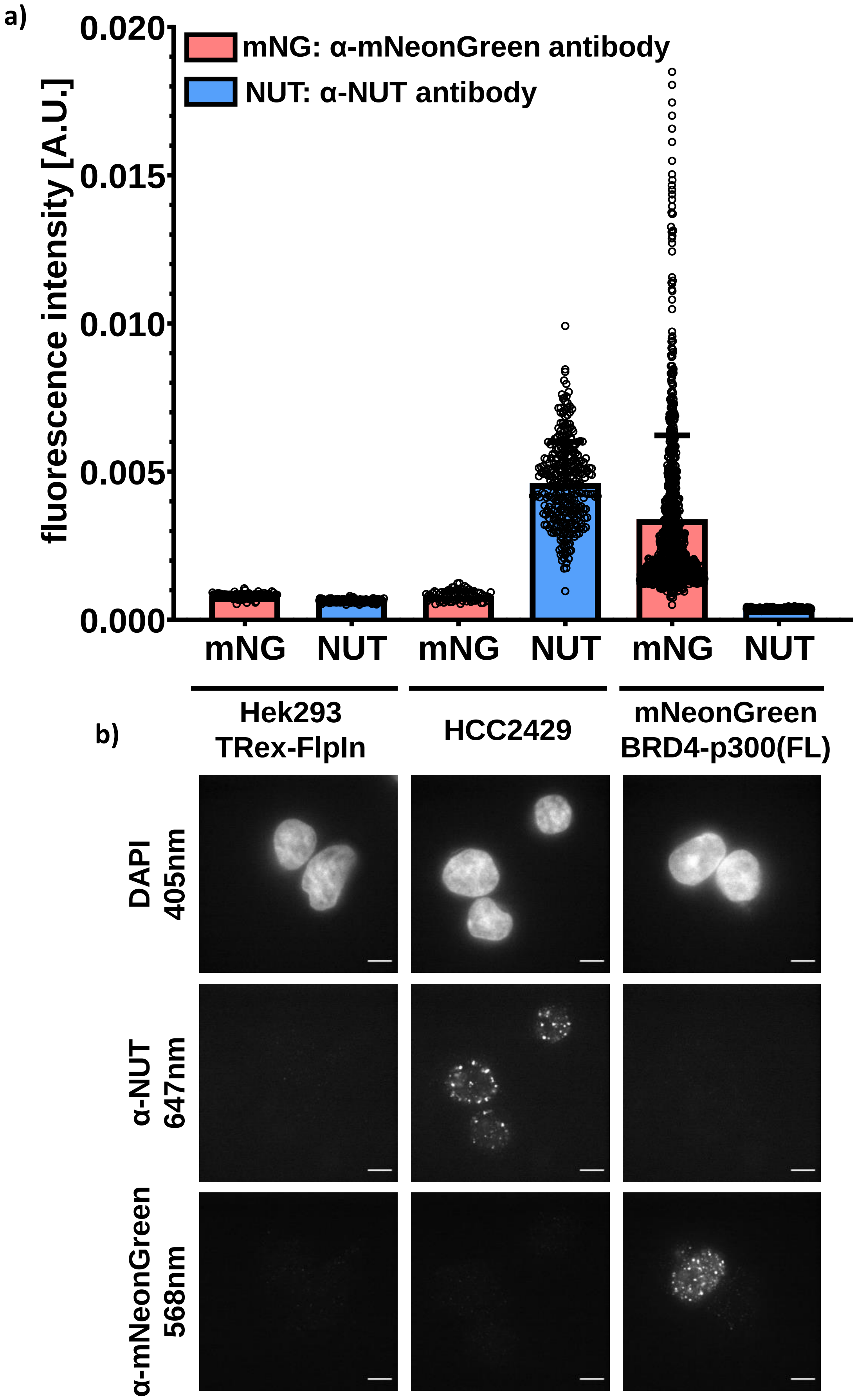

**Suppl. Fig. 7: Analysis of a potential bleed through between laser channels and cross-reactivity between the α-NUT and α-mNeonGreen antibodies.**

a) Average fluorescence intensity from α-NUT and α-mNeonGreen antibodies staining across different cell lines. Each datapoint is a single cell nucleus.

b) Representative micrographs of all three cell lines used in the analysis, as described. Scale bar = 10 µm.
